# Regulatory Rewiring Drives Intraspecies Competition in *Bacillus subtilis*

**DOI:** 10.1101/2025.11.27.690801

**Authors:** Margarita Kalamara, Alistair Bonsall, Jonathan Griffin, Joana Carneiro, Marek Gierlinski, Lukas Eigentler, David Stevenson, Amy Wood, Michael Porter, Helge C. Dorfmueller, Cait E. MacPhee, James C. Abbott, Nicola R. Stanley-Wall

## Abstract

Intraspecies interactions shapes microbial community structure and evolution, yet the mechanisms determining competitive outcomes among closely related strains remain unclear. The soil bacterium *Bacillus subtilis* is a model for microbial social interactions, where quorum-sensing systems regulate cooperation and antagonism. Here, we take a multifaceted approach to dissect the role of quorum-sensing regulation in competitive fitness. Isolate NCIB 3610 carries a signal unresponsive RapP-PhrP module that alters quorum-sensing control and promotes faster growth. Modelling and mutant analysis demonstrate that the small differences in growth rate conferred by RapP-PhrP^3610^ are sufficient to drive competitive exclusion. The importance of quorum sensing control is further exemplified by experimental evolution of distinct wild isolates, which revealed recurrent mutations in the sensor kinase *comP*, which phenocopy complete *comP* or *comA* deletions and confer a growth-linked competitive advantage. Key quorum sensing mechanisms are abandoned even in structured microbial communities, where it might be expected that communal traits are favoured. Furthermore, a phylogenomic survey of 370 *B. subtilis* genomes identified disruptive *comP* mutations in ∼16% of isolates. However, growth rate alone does not explain all interaction outcomes as even isogenic strains with equivalent doubling times differ in competitiveness. Transcriptomic profiling and validation experiments implicated a type VII secretion system toxin as an additional effector. These findings reveal that disruption of quorum-sensing pathways, whether naturally or through selection, provides a rapid route to competitive advantage, highlighting a fundamental trade-off between communal signalling and individual fitness in microbial populations.

**Importance:** Microbial competition and cooperation are key in shaping the structure and evolution of microbial communities. Our study on *Bacillus subtilis*, a model for microbial social interactions, reveals how alterations in cell-cell communication can enhance competitive fitness. We show, through a combination of modelling, mutational analysis, and experimental evolution, that certain strains of *B. subtilis* gain competitive advantage by disrupting their quorum sensing pathways, which leads to faster growth and enhanced competitiveness. Such mutations are prevalent in ∼16% of analysed genomes, underscoring their widespread evolutionary benefit under competitive conditions. Our research highlights the delicate balance between individual success and community cooperation in microbes, providing insights into microbial behaviour and potential applications in modifying microbial ecosystems.

## Introduction

Microbial life is inherently social. In nutrient-limited environments, microbes compete for survival but also cooperate through behaviours such as public goods sharing, division of labour, and collective defence actions. These interactions profoundly shape community structure and consequently ecosystem function, influencing not only bacterial populations but also their interactions with fungi, plants, and animals (1–3). Competition is particularly intense among closely related strains that share ecological niches, where outcomes depend on both cooperative investments and antagonistic strategies.

The Gram-positive bacterium *Bacillus subtilis* is an established model for dissecting microbial sociality (4). Within clonal biofilms, genetically identical cells differentiate into specialised subpopulations, coordinating cooperative matrix production (5) while also deploying antagonistic toxins to regulate growth and sporulation (6–9). In natural environments, however, *B. subtilis* strains frequently encounter conspecific rivals.

Such encounters can be shaped by kin discrimination, where strains cooperate with close relatives but compete with more distant lineages (10–12). Although molecules such as polymorphic toxins and secondary metabolites have been implicated in competitive outcomes (10–13), strain dominance cannot always be predicted from genomic content alone (14).

Central to *B. subtilis* social regulation are quorum-sensing systems that modulate cooperative and competitive behaviours. The ComQXPA system (Figure 1A) exhibits extensive polymorphism, partitioning isolates into distinct “pherotypes” that can communicate privately (15, 16). Parallel Rap–Phr modules further regulate ComA activity antagonistically (17, 18), and diversify social traits displayed by different isolates (19). Of intense study is strain NCIB 3610, which harbours an atypical RapP–PhrP system that is insensitive to its cognate peptide quorum sensing signal (20, 21) (Figure 1A). This variant alters the balance of cooperative regulation and tips the scales towards enhanced growth (22).

**Figure 1.**
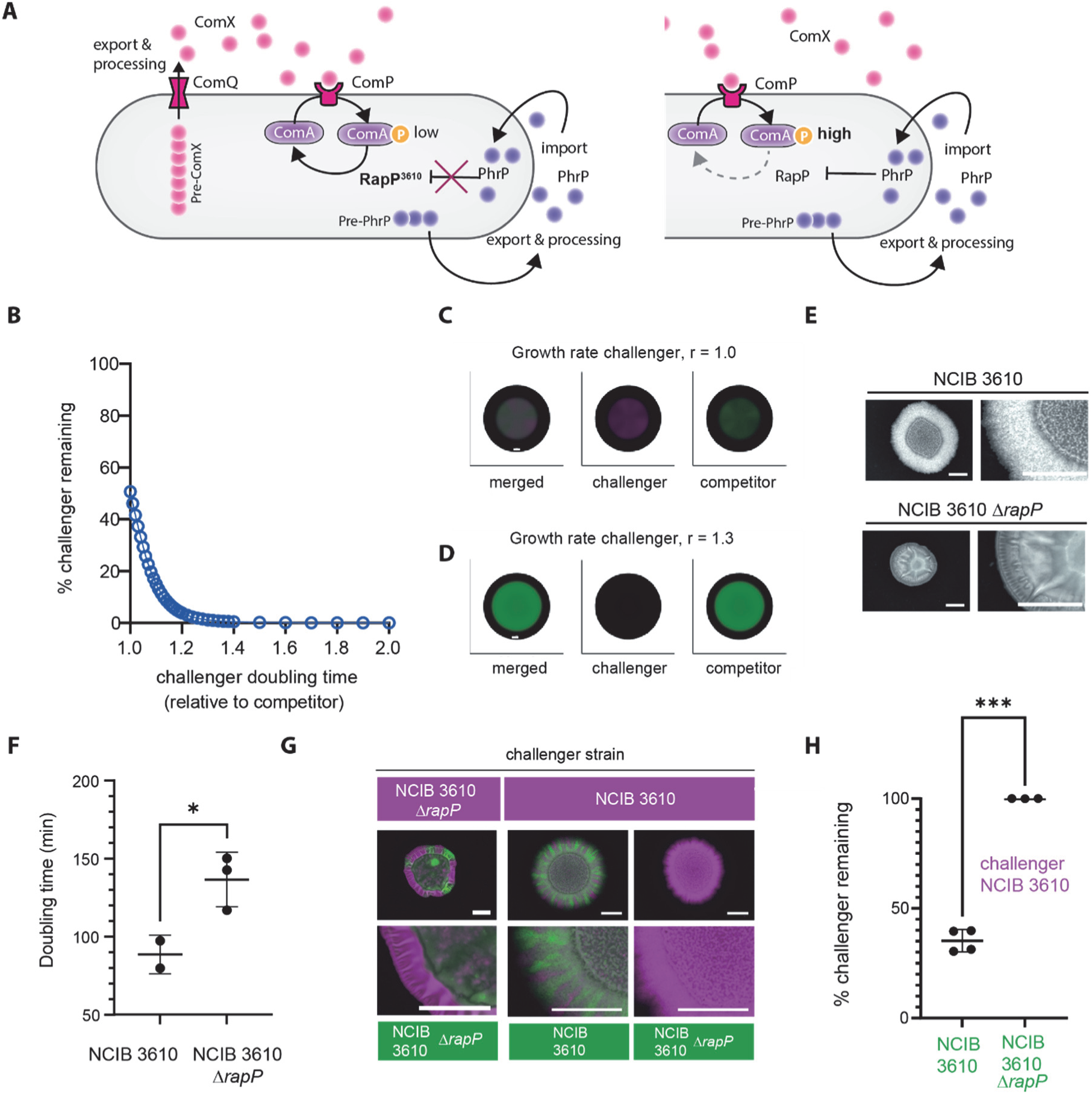
Deletion of *rapP* in NCIB 3610 decreases competitive fitness in mixed biofilms. (A) Schematic depicting the interconnections between the ComQXPA quorum sensing system and the RapP-PhrP quorum sensing module in NCIB 3610 (left) and other wild type isolates that contain a signal responsive RapP (right). ComX and PhrP are produced as pre-peptides that are exported and processed into the extracellular environment. At high cell density ComX binds ComP and triggers phosphorylation of the cognate response regulator ComA, while PhrP is imported to the cytoplasm. In NCIB 3610 RapP carries a single amino acid substitution (N236T) which renders it insensitive to PhrP levels and thus the level of phosphorylated ComA remains low due to the dephosphorylation activity of RapP. In either a *rapP* deletion strain (not depicted) or in isolates where RapP is responsive to PhrP the level of phosphorylated ComA can rise in line with increases in cell density. (B) Relationship between competitive outcome and relative growth rates of *in silico* strains *B1* and *B2*. (C-D) Model simulations with equal growth rates (C) or with *B2* having a growth advantage (relative growth rate = 1.3) (D). Black areas show the computational domain (Ω). Scale bars represent 28 nondimensional units. Strains are shown individually (*B1*, green; *B2*, magenta) and merged (grey = overlap). (E) Colony biofilms of NCIB 3610 and NCIB 3610 Δ*rapP* grown for 48 h at 30°C. Scale bars, 0.5 cm. The Δ*rapP* strain expressed mTagBFP (NRS7280). (F) Doubling time (min) during exponential growth at 30°C with shaking. *p* ≤ 0.05, unpaired *t* test with Welch’s correction. Each point = one sample; error bars = SD. (G) Representative dual-strain colony biofilm after 48 h at 30°C. Fluorescence signals from mTagBFP (magenta) and GFP (green). Scale bar, 10 mm. (H) Quantification of competitive outcomes as % challenger NCIB 3610 (mTagBFP+) remaining providing a measure of relative strain density. Each point = one colony biofilm; error bars = SD. ****p < 0.0001, unpaired *t* test with Welch’s correction.

The mechanisms that confer competitive superiority to NCIB 3610, and whether susceptible strains can evolve counter-adaptations, remain poorly understood. Here, we combine experimental evolution, genetic reconstruction, transcriptomics, and comparative genomics to dissect the role of quorum-sensing regulation in competitive fitness. Our results reveal that the presence of the signal unresponsive RapP-PhrP module, or disruption of the ComQXPA system, underpin the strain’s ability to dominate diverse environmental isolates in colony biofilm competitions. Furthermore, disruption of the ComQXPA systems provides a recurrent evolutionary strategy both in laboratory and natural populations that uncouples cells from costly regulation, accelerates growth, and enhances competitiveness in environments dominated by conspecific rivals. However, intraspecies interactions cannot be reduced to growth rate alone: here we demonstrate that isogenic mutants with equivalent growth diverge in competitive ability. Transcriptomic profiling implicates toxins, including those secreted by type VII secretion systems, as additional drivers of these outcomes.

## Results

### Signal unresponsive RapP of *B. subtilis* NCIB 3610 impacts pairwise intraspecies competition outcomes

We previously demonstrated that *B. subtilis* strain NCIB 3610 dominates pairwise interactions when co-cultured with an array of *B. subtilis* isolates in a dual isolate colony biofilm setting (14). *B. subtilis* quorum sensing is regulated by ComQXPA and a suite of Rap-Phr accessory systems (Figure 1A). The allele of RapP encoded by NCIB 3610 is associated with increased fitness in an otherwise isogenic context (23) and linked with an accelerated growth rate (22, 23). We therefore questioned whether it is NCIB 3610 RapP-PhrP (RapP-PhrP^3610^) that drives the successful outcome for NCIB 3610 during intraspecies interactions. We first tested this hypothesis by adapting an existing mathematical model (24) which describes the spatio-temporal dynamics of two isolates in a single growing biofilm. For simplicity, the partial differential equations model reduces the dynamics to local growth through a logistic growth function and spatial spread through negative density-dependent diffusion (see Methods). Our *in-silico* modelling revealed that a seemingly small difference in doubling times of isolates co-cultured in a single biofilm manifested in a large impact on the competition outcome. The *in-silico* isolate with the shorter doubling time rapidly dominated the pairwise interaction (Figure 1B-D; (24)).

To explore the role of RapP^3610^ in mediating the competitiveness of NCIB 3610, we constructed an NCIB 3610 Δ*rapP* mutant and noted a mucoid colony biofilm morphology (Figure 1E) (21), reminiscent of that formed by environmental isolates of *B. subtilis* (Figure S1). Measuring the doubling time under planktonic growth conditions during exponential phase growth revealed that NCIB 3610 was on average ∼1.4-fold faster compared to the otherwise isogenic NCIB 3610 Δ*rapP* mutant (Figure 1F). Consistent with growth contributing to the competition outcome, NCIB 3610 readily outcompeted NCIB 3610 Δ*rapP* in a dual isolate colony biofilm competition setting (Figure 1G-H). In these experiments, we employed NCIB 3610 Δ*rapP* expressing mTagBFP (NRS7280) and co-cultured it with NCIB 3610 constitutively expressing GFP (NRS6942). As control experiments, we co-cultured isogenic variants of NCIB 3610 and NCIB 3610 Δ*rapP* which differed in whether they produced GFP or mTagBFP as the “reporter”. As anticipated, each of the control strain combinations showed coexistence in the mature population (Figure 1G-H).

### The impact of RapP on the competition outcome for an array of *B*. *subtilis* isolates

To delve deeper into the role of RapP in mediating the competitiveness of NCIB 3610 we examined pairwise interactions between the NCIB 3610 Δ*rapP* mutant (constitutively producing mTagBFP (NRS7280)) and a suite of different wild-type *B. subtilis* isolates, modified to constitutively produce GFP, that we had previously used (14). We overlayed the data obtained with our previously published results that used NCIB 3610 (mTagBFP+) as the challenging strain (14) (Figure 2A) and calculated the impact of RapP on the outcome of each pairwise interaction (Figure 2B). Our findings demonstrate a strain-dependent contribution of RapP to NCIB 3610’s highly competitive phenotype. Some strains that could co-exist with NCIB 3610, albeit in minimal proportions, became more prevalent in the mature colony biofilm when cocultured with NCIB 3610 Δ*rapP*. These strains included NRS6202, NRS6105, NRS6145, NRS6107, NRS6116, NRS6118, NRS6121, and NRS6190. Other strains that were entirely outcompeted by NCIB 3610 remained virtually undetectable within the population (e.g., NRS6103, NRS6132, NRS6181, and NRS6187). By aligning the interaction outcome with strain phylogeny, we established that the RapP-mediated contribution to NCIB 3610 competitiveness was broadly associated with the relatedness of the soil isolate to NCIB 3610 (Figure 2C). The more closely related the co-cultured strain was to NCIB 3610 in terms of phylogeny, the greater the extent of RapP’s impact compared with more distantly related isolates.

**Figure 2.**
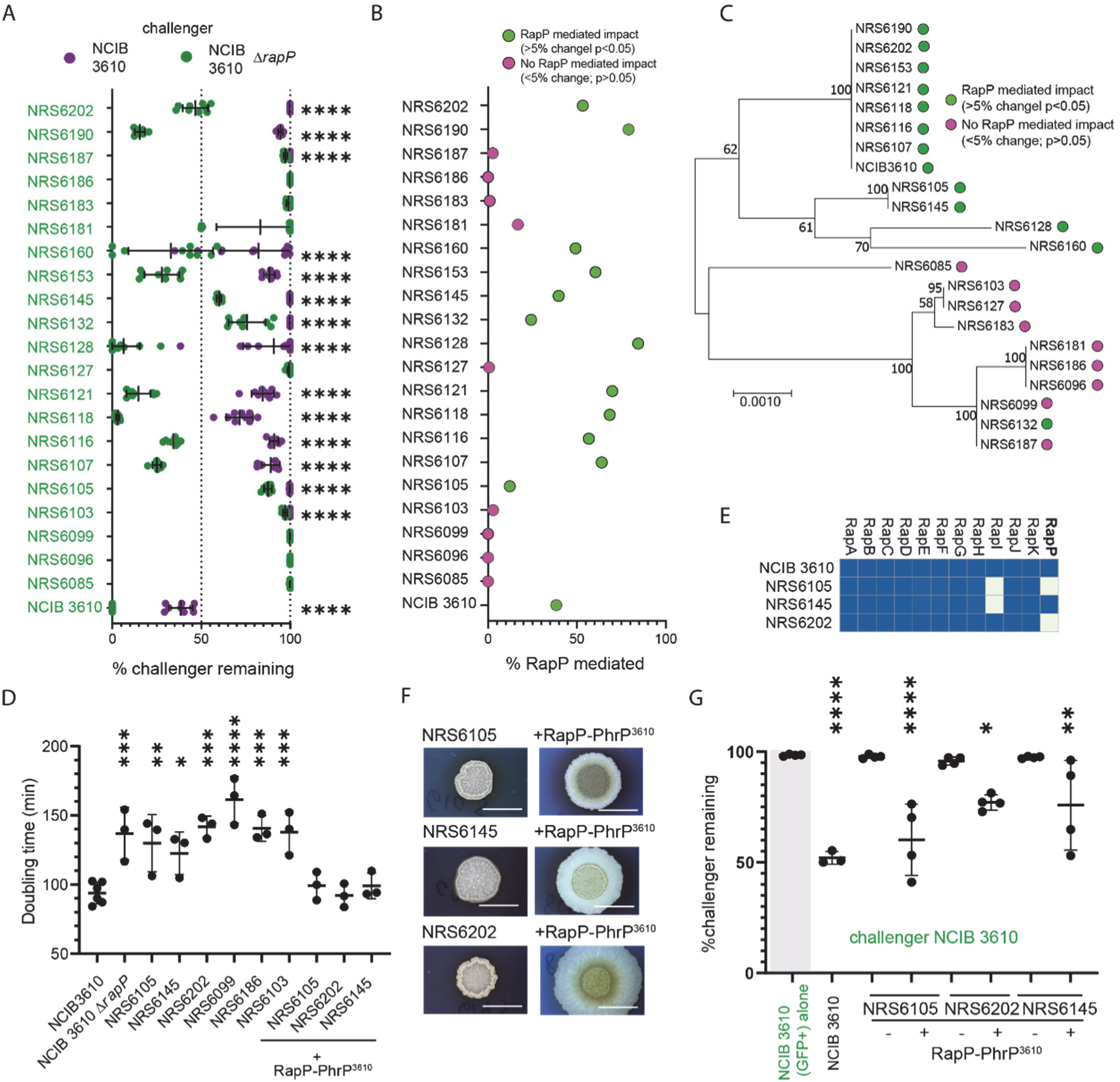
The impact of RapP on NCIB 3610 competitive fitness is isolate dependent. (A) Quantification of pairwise biofilm competitions shown as % challenger remaining (relative strain density). Challengers were NCIB 3610 (mTagBFP+, magenta; data from (14)) or NCIB 3610 *ΔrapP* (mTagBFP+, green). Partner strains are indicated on the y-axis. Each point = one biofilm; error bars=SD. p < 0.0001, unpaired t test. Non-significant comparisons not shown. (B) RapP effect on competition outcomes, calculated as (% remaining after Δ*rapP* co-culture – % remaining after NCIB 3610 co-culture). The definition of a RapP effect is designated in the legend and was >5% change in the interaction outcome and a p value of <0.05. (C) Maximum-likelihood phylogeny based on *gyrA, rpoB, dnaJ,* and *recA*, with bootstrap values shown as branch labels (adapted from (14)). Strains showing a RapP effect are marked as in (B). (D) Doubling time (min) of each strain during exponential growth at 30 °C with shaking. a p-value of ≤0.05 (*); ≤0.01 (**); ≤0.001 (***) or ≤0.0001 (****) vs. NCIB 3610 (one-way ANOVA, Dunnett’s test). Each point = one sample; error bars = SD (E) Presence/absence of *rap* genes in NCIB 3610, NRS6105, NRS6145, and NRS6202. (F) Representative colony biofilms grown for 48 h on biofilm-inducing medium at 30°C. Scale bars, 10 mm. (G) Quantification of % NCIB 3610 (gfp+) remaining after co-culture with the indicated strain for 48 h. “–ve” = NCIB 3610 alone; “+ve” = NCIB 3610 (gfp+) (NRS6942) alone. RapP-PhrP presence (+) or absence (–) in the chromosome is shown in the legend. Each point = one biofilm; error bars = SD. p thresholds as in (D).

To explore if the variable impact of *rapP* on the competition outcome was underpinned by differences in growth, we determined the doubling time during the exponential phase of planktonic growth for a selection of the strains (Figure 2D). We chose three strains that showed a competitive advantage against NCIB 3610 Δ*rapP* (NRS6105, NRS6145, NRS6202), and three strains (NRS6103, NRS6099, NRS6186) that showed no change in the interaction outcome. We determined that each strain had a significantly slower doubling time compared to NCIB 3610 but comparable to that of NCIB 3610 Δ*rapP*. Therefore, growth rate is not the sole determinant of the interaction outcome. This conclusion is supported by the outcome of the pairwise interaction screen when the closely phylogenetically related isolate NRS6202 replaced NCIB 3610 / NCIB 3610 Δ*rapP* as the challenger strain. The same pattern of interaction outcomes against the other isolates was measured as when NCIB 3610 Δ*rapP* was the challenger (compare Figure S2B and Figure 2A).

Rap-Phr protein pairs are found as discrete modules that are found in variable number in different wild isolates of *B. subtilis* (19). This allows for plasticity in quorum sensing responses. Analysis of 370 complete *Bacillus subtilis* genomes (see methods) detected the *rapP* coding region in 38 strains (41 genome sequences, accounting for duplicate sequencing of the same strain, e.g. NCIB 3610). 23 of these strains, which included NCIB 3610, carry *rapP* on a plasmid (Figure S2A). Seven of the 38 *rapP*-encoding strains harbour the NCIB 3610 variant of the *rapP* coding region that is PhrP unresponsive. Metabolic activity can be rewired when RapP^3610^ is transferred to a closely related domesticated strain of *B. subtilis* (22). We sought to understand if introduction of the RapP-PhrP^3610^ coding region into more distantly related wild-type isolates of *B. subtilis* would also accelerate growth and correspondingly enhance competitiveness against NCIB 3610. In essence, this represents the reciprocal of deleting *rapP* from NCIB 3610 and observing a drop in competitiveness. We used NRS6105, NRS6145, and NRS6202, of which, NRS6145 encodes RapP-PhrP (the signal-responsive variant) (see Supplemental File F1) (Figure 2E). We introduced the *rapP-phrP^3610^* coding region under the control of the native NCIB 3610 *rapP* promoter at the ectopic *amyE* location on the chromosome.

Introduction of RapP-PhrP^3610^ conferred an “NCIB 3610-like” colony biofilm morphology in each isolate (Figure 2F) and a decrease in the doubling time measured in planktonic cultures during exponential phase (Figure 2D). In addition, there was an increase in competitive fitness when co-cultured with NCIB 3610 (gfp+) compared to the respective parental strains. For these dual isolate co-cultures, the proportion of NCIB 3610 (gfp+) remaining in the mature population was measured using flow cytometry (Figure 2G). Collectively, these data demonstrate that RapP-PhrP^3610^ is dominant over the native RapP-PhrP encoded by NRS6145, and that introduction of RapP-PhrP^3610^ increases an isolate’s ability to endure an encounter with NCIB 3610.

### Experimental evolution of *B. subtilis* soil isolates

Given the rarity of *rapP* (Figure S2) and the knowledge that the genetic landscape of strains can adapt under stress, we aimed to identify other spontaneous genetic adaptations that could enhance the ability of *Bacillus subtilis* strains (normally outcompeted by NCIB 3610) to withstand such encounters. We selected NRS6105, NRS6145, and NRS6202 as each of these strains is largely outcompeted by NCIB 3610 within a pairwise interaction in a dual isolate colony biofilm competition setting (Figure 3A), but the isolates can persist at a low level (for example, Figure 3B), and each can be re-isolated as single colonies from the mixed population (Figure 3C).

**Figure 3.**
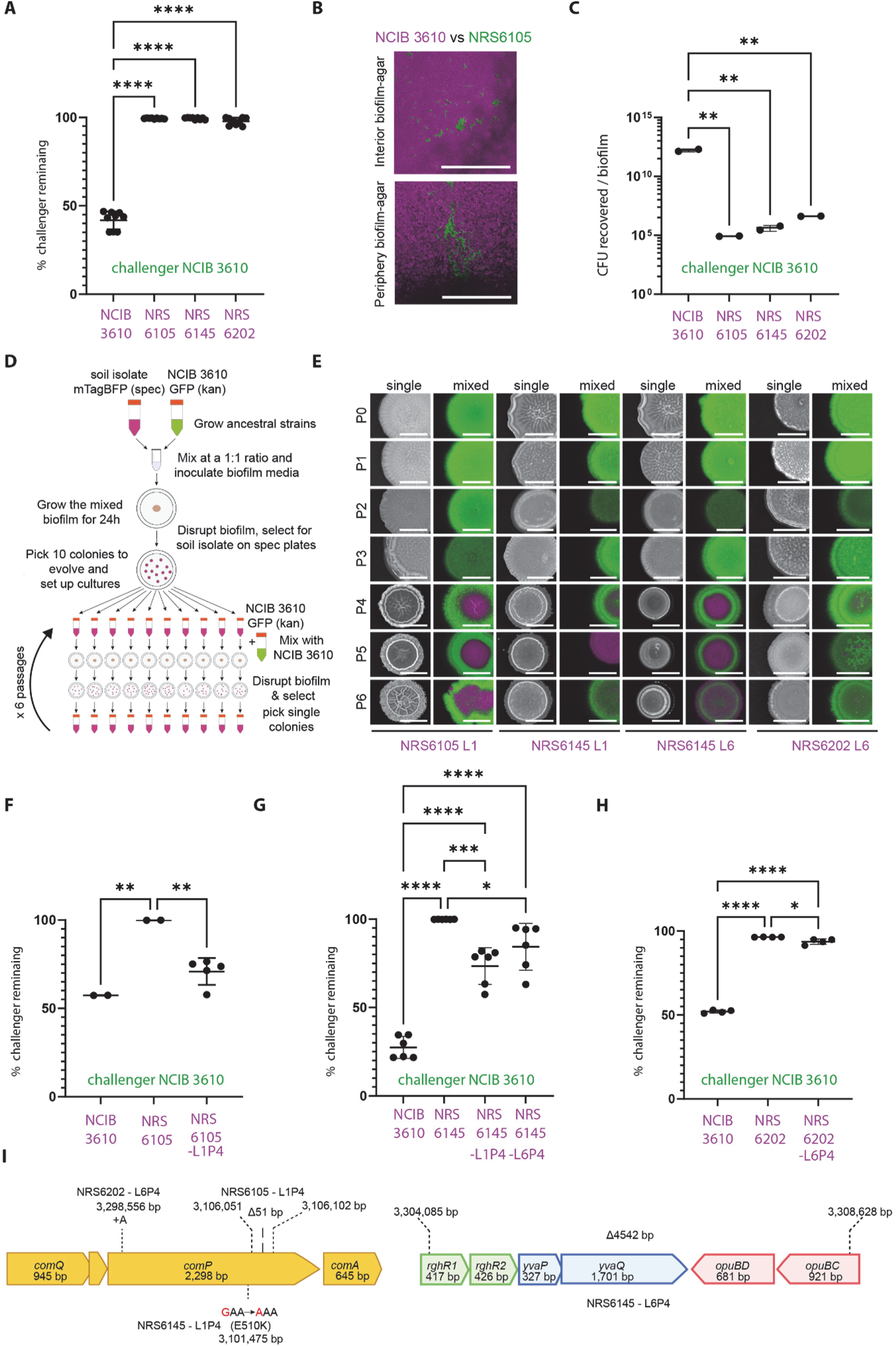
Experimental evolution in dual-isolate colony biofilms. (A) Prevalence of NCIB 3610 (gfp+) after 24 h in dual biofilms, replotted from (14). Data shown as % challenger remaining (relative strain density). (B) Representative confocal image of NCIB 3610 (magenta) with NRS6105 (green) after 24 h. Scale bar, 500 μm. (C) Colony-forming units (CFU) per biofilm after 24 h co-culture at 30 °C. p-value of ≤0.05 (*); ≤0.01 (**); ≤0.001 (***) or ≤0.0001 (****), one-way ANOVA with Tukey’s test. Non-significant comparisons not shown. (D) Experimental design schematic: soil isolates (NRS6105, NRS6145, NRS6202) co-cultured with NCIB 3610; evolved clones re-isolated on spectinomycin plates. (E) Colony biofilms of ancestral and evolved isolates grown alone (“single”) or mixed with NCIB 3610 (gfp+) for 24 h. NCIB 3610 = green; soil isolates = magenta. P1–P6 denote passages; strain background and lineage indicated below. Scale bars, 3 mm. (F–H) Quantification of the prevalence of NCIB 3610 after 48 h in co-culture with ancestral and evolved lineages as measured by relative strain density: (F) NRS6105, (G) NRS6145, (H) NRS6202. Statistics as in (C). (I) Schematic of mutations identified in evolved isolates. Dashed lines and numbers indicate genomic positions; strain backgrounds are indicated.

We used derivatives of the NRS6105, NRS6145, and NRS6202 that were engineered to be spectinomycin resistant and constitutively produce mTagBFP (the derived isolates are called NRS6936, NRS6951, and NRS7201 respectively and will be referred to for simplicity by the parent strain name (mTagBFP+)). We used a variant of NCIB 3610 that constitutively expressed GFP and was resistant to kanamycin as the challenger strain (NRS1473, NCIB 3610 (gfp+)) (See Supplemental File F2, Table S1). We conducted 6 passages in which NRS6105 (mTagBFP+), NRS6145 (mTagBFP+), and NRS6202 (mTagBFP+) were co-incubated with NCIB 3610 (gfp+) in mixed isolate colony biofilms for 24h (Figure 3D). NRS6105 (mTagBFP+), NRS6145 (mTagBFP+), and NRS6202 (mTagBFP+) were selectively recovered each time using spectinomycin antibiotic resistance selection. We conducted this experiment concurrently for 10 lineages of each soil isolate (three of the lineages of NRS6202 (mTagBFP+) were stopped prematurely) (Figure 3E, Figure S3A).

Of the 27 lineages generated, 6 lineages became better able to co-exist with NCIB 3610 (gfp+). After validation, the number of lineages was reduced to four, which comprised two lineages derived from the NRS6145 (mTagBFP+) parental strain, one lineage from NRS6105 (mTagBFP+), and one from NRS6202 (mTagBFP+) (Figure 3E&S3A-C). In each case, the change in the ability to co-exist with NCIB 3610 (gfp+) became apparent from the 4^th^ passage (Figure 3E-H). For each of the evolved strains that displayed an enhanced ability to co-exist, the mixed isolate colony biofilm that formed showed a distinct spatial distribution of the two strains within the overall architecture when imaged (Figure 3E-H). The evolved soil isolates dominated the middle of the biofilm while NCIB 3610 (gfp+) occupied the periphery, essentially encircling the evolved isolates. In addition to the change in the phenotype of the mixed biofilms, the single isolate colony biofilms of the evolved variants showed a different phenotype to their parental isolates, such that they appeared less mucoid and occupied a reduced footprint (Figure S3D).

We acquired short-read genome sequencing data for each of the evolved isolates at passage 3 and 4. Sequence analysis revealed that in all four lineages, there was a single genetic change present between the passages. For three out of four evolved isolates, the mutations appeared within the *comP* coding region, a core component of the *B. subtilis* quorum sensing system (recall Figure 1A). There was one *comP* mutant from each of the strain backgrounds = subjected to evolution. The *comP* coding region of the evolved isolate of NRS6202 contained a single adenine insertion at chromosome position 3,298,556 bp which resulted in a truncation of the *comP* coding region. In strain NRS6105 the nucleotides that encode amino acids 633-649 (inclusive) of ComP were excised. These 17 amino acids reside in a helical structure in proximity to the catalytic and ATP binding domain, specifically in helix α8 (Figure SE-F). Although this deletion does not directly interfere with ATP binding, based on structural modelling (Figure SG-H), and the side chain of His691 remains in proximity to the ATP binding site, it likely alters the overall conformation of the catalytic domain and its spatial relationship within the ComP dimer, potentially affecting the interface with the cognate response regulator and substrate ComA. Therefore, ComPΔ633-649 variant is expected to be defective of phosphorylating its cognate response regulator ComA. In the evolved isolate of NRS6145 (L1P4) the chromosome contained a single nucleotide mutation at position 3,101,476 bp which yielded an amino acid substitution in ComP - E510K. We again generated structural models for both the ComP^NRS6145^ (WT) and ComP^NRS6145_E510K^ mutant sequences using AlphaFold3 (25) (see methods). In the WT model, a potential hydrogen bond is observed between the side chain of E510 and R331, with both residues being highly conserved among ComP homologues (See Supplemental File F3). Additionally, E394, located on an adjacent α-helix, is positioned within hydrogen-bonding distance to R331, suggesting a stabilising three-residue interaction network that may contribute to the relative packing of these helices. In contrast, inspection of the E510K mutant models indicates a reorganisation of a hydrogen-bonding network.

The lysine side chain at position 510 does not maintain the interaction geometry with R331 as seen for wild type ComP^NRS6145^. Instead, K510 can form a hydrogen bond with nearby residues (e.g., E394), while R331 appears displaced, and its distance to residue 510 exceeds typical hydrogen bond cutoffs. This rearrangement suggests that the local tertiary structure of the three helices may be altered, effectively “locking” the region into an alternative conformation compared to the WT protein.

Taken together, the modelling results point to a rewiring of hydrogen-bonding interactions between residues E/K510, R331, and E394, consistent with the observed functional loss of activity. The last mutant analysed was NRS6145-L6P4 which contained a 4,542 bp deletion in the chromosome. The region deleted covers the majority of the *rhgR1* and *opuBC* genes along with all the genes between them (*rghR2*, *yvaP*, *yvaQ,* and *opuBD*) (Figure 3I) (see Supplemental File F1). Given that three of the four mutations mapped to *comP*, subsequent analyses focused on validating the role of *comP* disruption in determining the competitive fitness of *B. subtilis* isolates during intra-species interactions in a dual-isolate colony biofilm model.

### Impacts associated with mutation of the ComQXPA quorum sensing system

To validate the role of ComP in shaping intra-species interactions we reconstructed the *comP*Δ633–649 deletion (Figure 4A-B) and a complete in-frame *comP* deletion in isolate NRS6105. Each *comP* mutant strain (mTagBFP+) was challenged against NCIB 3610 (GFP+) in dual-isolate colony biofilms. The outcomes of these competitions resembled those observed during our experimental evolution experiments (compare Figure 3 with Figure 4A–C). Both *comP* mutants were more prevalent than the parental NRS6105 strain, although NCIB 3610 maintained a competitive advantage. These results show that deletion of residues 633–649 produces a phenotype equivalent to a full *comP* deletion (Figure 4A–B), revealing that, consistent with the *in silico* structural analysis (Figure S3G-H, ComP is inactivated.

**Figure 4.**
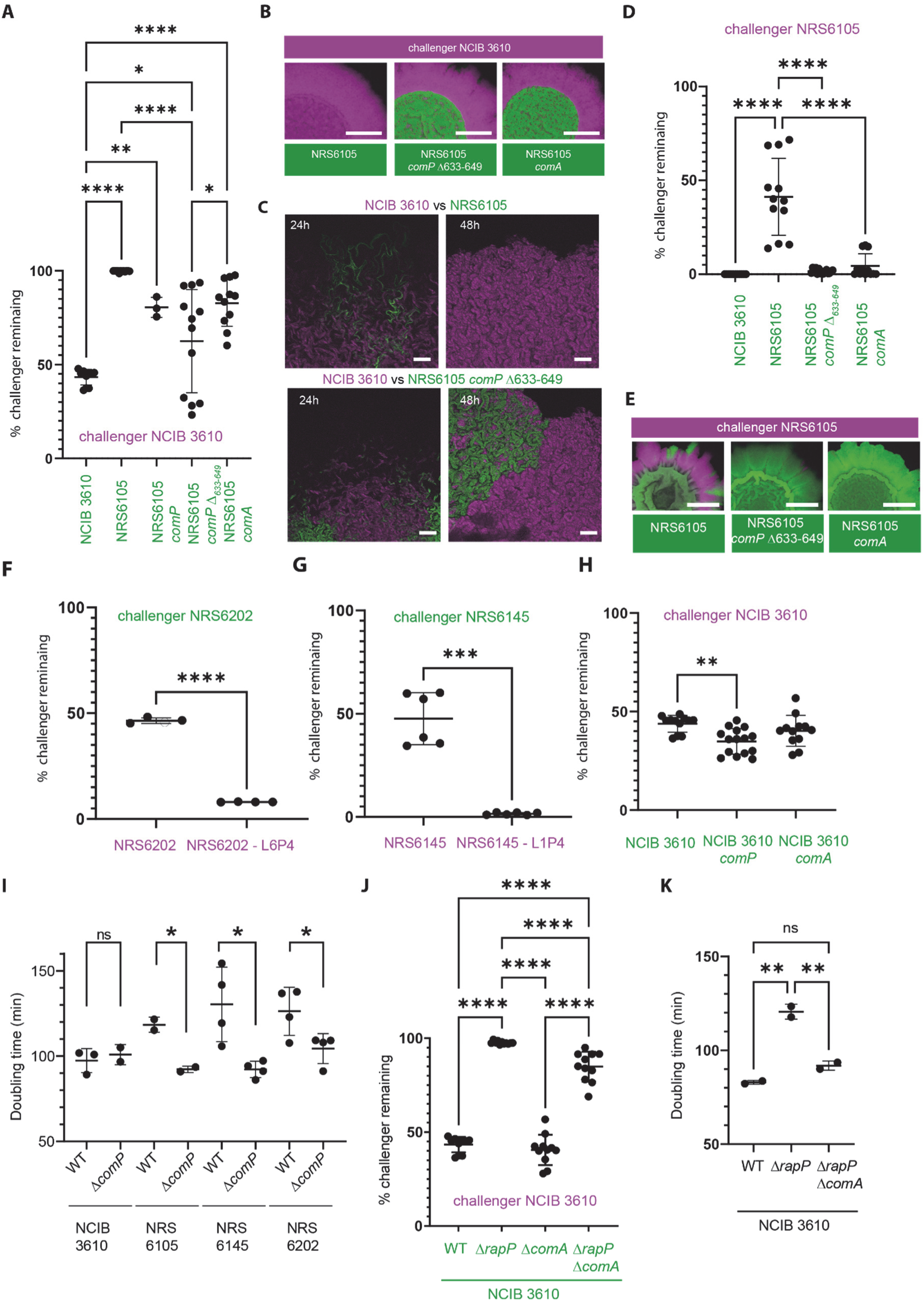
ComQXPA influences the outcome of intraspecies biofilm interactions. (A) Quantification of NCIB 3610 (gfp+) prevalence (relative density) after 48 h in dual colony biofilms with partner strains (mTagBFP+). % challenger remaining was measured by image analysis. **p < 0.0001, one-way ANOVA with Sidak’s test. (B) Representative colony biofilms corresponding to (A). Scale bars, 10 mm (left) or 3 mm (zoom). (C) Confocal images of NCIB 3610 (gfp+) vs. NRS6105 or NRS6105 Δ*comP* (mTagBFP+) at 24 h and 48 h. Scale bar, 100 μm. (D) Prevalence of NRS6105 (gfp+) in dual biofilms with indicated partner strains after 48 h. (E) Representative images for (D). Scale bars, 10 mm (left) or 3 mm (zoom). (F–H) Outcomes of dual biofilms with additional isolates. % challenger remaining was measured by image analysis. Significance determined by Welch’s t test (F–G) or ANOVA with Tukey’s test (H). Non-significant comparisons not shown. (I) Doubling times of NCIB 3610 and indicated isolates during exponential growth at 30 °C. p ≤ 0.05, Welch’s t test. (J) Interaction outcomes with NCIB 3610 (mTagBFP+) as challenger and the indicated strains (gfp+). ****p < 0.0001, ANOVA with Tukey’s test. (K) Doubling times of NRS6942, NRS7279, and NRS7771 at 30 °C. p ≤ 0.01, ANOVA with Tukey’s test. Note - Data for NCIB 3610 in Figure 4A, 4J and 4H & NCIB 3610 Δ*comA* in Figure 4J and 4H are the same dataset.

We next assessed whether the impact of deleting *comP* in NRS6105 was mediated through its cognate response regulator ComA. To test this, we generated an NRS6105 Δ*comA* mutant (GFP+) and challenged it against NCIB 3610 (mTagBFP+) in dual-isolate biofilms. Like the *comP* mutant, the Δ*comA* strain displayed increased survival compared to the parental NRS6105 isolate (Figure 4A). These findings suggest that disruption of the ComP–ComA regulatory axis enhances competitiveness of NRS6105 in mixed biofilms. This phenotype mirrors the case in NCIB 3610, where deletion of *rapP* elevates ComA∼P levels and reduces its competitiveness (20, 22).

To assess whether the competitive advantage conferred by NRS6105 *comP* and *comA* mutations was specific to interactions with NCIB 3610, we also competed these mutants against their parental NRS6105 background. In both cases, the mutants strongly outcompeted the parental strain, occupying nearly the entire colony biofilm when starting from a 1:1 inoculum (Figure 4D–E). Similar results were obtained for evolved isolates: *comP* mutants of NRS6145 and NRS6202 dominated their respective parental strains (Figure 4F–G). In contrast, *comP* or *comA* mutations introduced into NCIB 3610 itself produced only modest changes when competed against the NCIB 3610 parent consistent with the presence of RapP^3610^ (Figure 4H).

Like RapP-PhrP^3610^ (22), ComQXPA is linked to a trade-off in growth and antibiotic production (26). We next examined whether disruption of the ComQXPA system influences growth rate. Deletion of *comP* or *comA* reduced doubling times in all backgrounds tested except NCIB 3610 (Figure 4I). For NRS6105 Δ*comP*, as well as evolved NRS6145 L1P4 and NRS6202 L6P4 strains, growth rates during exponential phase approached those of NCIB 3610 and differed significantly from their parental isolates. These findings are consistent with unequal growth being a major driver of exclusion of parental strains by their *comP*-mutant derivatives, and with accelerated growth underpinning the ability of these mutants to coexist with NCIB 3610.

Finally, to complete our interrogation, we investigated the relationship between ComA and RapP in NCIB 3610. Since NCIB 3610 already exhibits low ComA∼P activity due to RapP-PhrP^3610^ (20) we hypothesised that *comA* would act epistatically to *rapP*. To test this, we constructed an NCIB 3610 Δ*rapP* Δ*comA* double mutant and compared its biofilm phenotype with the single mutants. The double mutant more closely resembled the Δ*comA* strain in biofilm morphology and the surface area (footprint) occupied by the biofilm (Figure S4D). In competition assays, the double mutant displayed enhanced fitness relative to Δ*rapP* alone but did not reach the coexistence level observed for Δ*comA* (Figure 4J). Notably, this occurred despite recovery of the doubling time shown by the Δ*comA* mutant to wild-type levels (Figure 4K). These results suggest that RapP’s impact on NCIB 3610 competitiveness is only partially mediated through ComA, and that equalising growth rates alone does not guarantee coexistence in colony biofilms.

### Towards identification of specific competition determinants

To explore the molecular basis of NCIB 3610’s competitive advantage, we performed RNA sequencing to obtain the global transcriptional profiles of NCIB 3610 (gfp+) and its Δ*rapP* Δ*comA* (gfp+) derivative. These strains were chosen because they show equivalent growth rates in planktonic culture, yet NCIB 3610 consistently outcompetes the Δ*rapP* Δ*comA* mutant in dual-isolate colony biofilms (see Figure 4). RNA was isolated from each strain after 20 hours of incubation under colony biofilm conditions and transcriptomic analysis revealed 469 differentially expressed transcripts between *B. subtilis* NCIB 3610 and its Δ*rapP* Δ*comA* mutant, as defined by FDR < 0.01 and |logFC| > 1 (Figure S5A).

Despite the comparable growth rates in planktonic culture, KEGG pathway and GO-term analysis revealed down-regulation of growth and metabolism associated pathways in Δ*rapP* Δ*comA* (See Supplemental File F2, Table S2). This is consistent with the rewiring of metabolism that occurs when *rapP*^3610^ is deleted (22). While metabolic rewiring may contribute to the interaction outcome, proteinaceous toxins and differential antibiotic expression have also been shown to influence intraspecies interactions. For example, toxins secreted by the Type VII secretion system (T7SS) influence competition among *B. subtilis* isolates grown in biofilms (13). Therefore, we examined expression of 89 known toxin and toxin-associated transcripts (See Supplemental File F2, Table S3).

We noted that transcripts from the *yfj, pks, ywq, alb,* and *wap* toxin-encoding operons were significantly down-regulated in the Δ*rapP* Δ*comA* strain (Table S3). Of these, the *yfj* operon has a minimal impact on intraspecies competition in biofilms (Kobayashi, 2023) and therefore was excluded from further analysis. The *pks* operon, controlling bacillaene synthesis, is linked to interspecies competition (Vargas-Bautista et al., 2014). The operon is present in all isolates in our analysis (14) and therefore its simple presence or absence does not correlate with the intraspecies outcomes we observe and was excluded. For the same reason (14), the *alb* operon, which contributes to the export of SboA, the bacteriocin subtilosin (Zheng et al., 1999) was excluded. The *wap* operon is polymorphic and contains the *wapA* toxin and *wapI* immunity genes. WapA mediates contact-dependent competition (Koskiniemi et al., 2013) but while the Δ*wapAI* mutant was outcompeted by the parental strain in liquid culture there was a lesser impact in biofilms (Kobayashi, 2023). This left the *ywq* operon (Figure 5A) which is linked with intraspecies competition mediated by the Type VII secretion system (13). The RNA sequencing analysis showed that expression of all members of the *ywq* operon were all downregulated over 2-fold (Figure 5B). In additional support of YwqJ having a role as an intraspecies competition determinant, our bioinformatic analysis of the T7SS toxin genes (which contain LXG-domains) carried by NCIB 3610 revealed considerable variation in their prevalence across the environmental isolates used in this study (Figure 5C; Figure S5B). More specifically, we noted that isolates that were better able to coexist with NCIB 3610 derived strains displayed greater similarity in their T7SS repertoires compared to isolates that were excluded.

**Figure 5:**
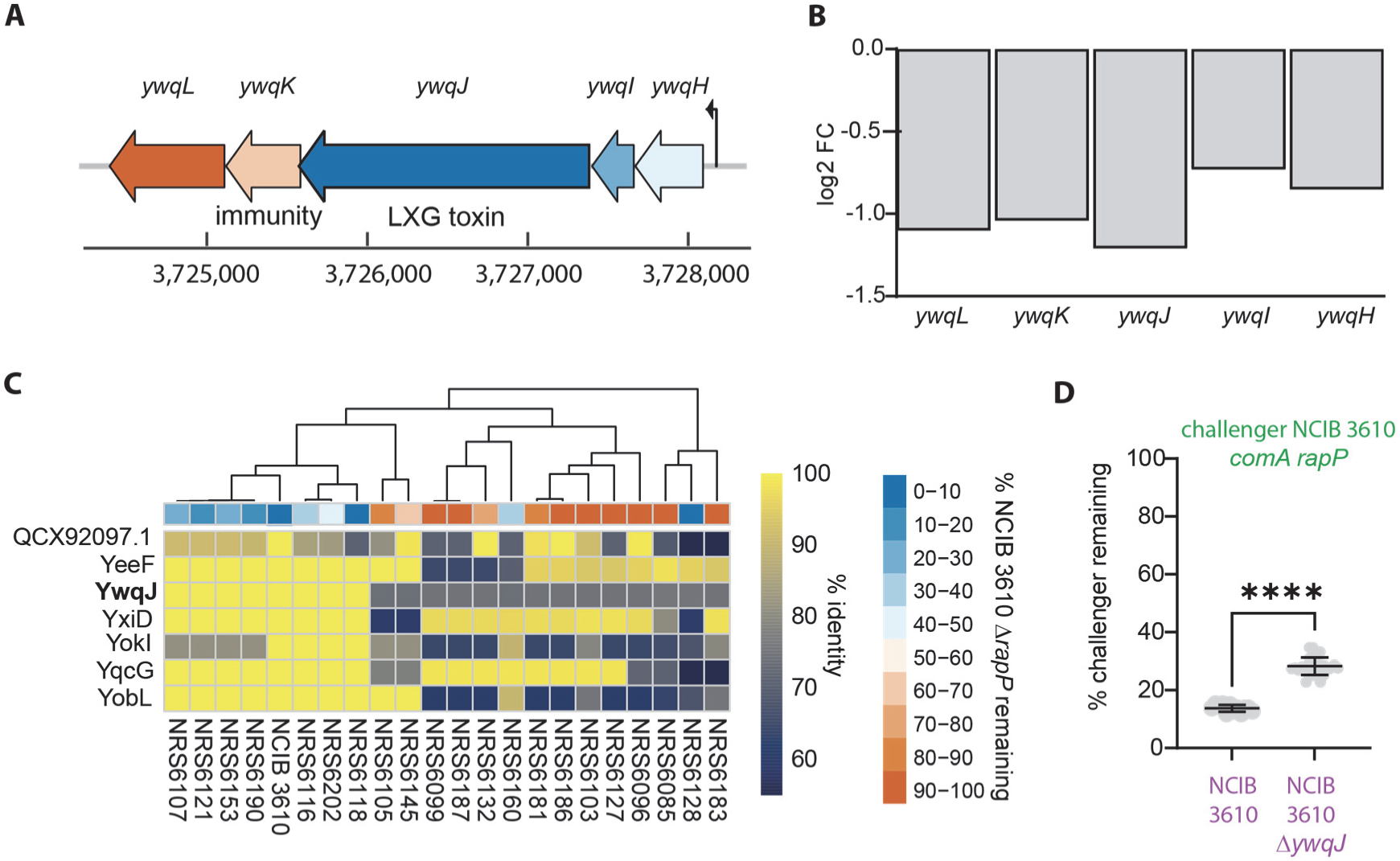
The Type VII section system toxin YwqJ is a key determinant. (A) Schematic of the *ywq* operon based on genomic information extracted from Subtiwiki (27). (B) RNA sequencing log2FC values for the members of the *ywq* operon. (C) Heatmap depicting the % protein sequence identity of the top matched LXG containing protein from the suite of *B. subtilis* isolates (x-axis) relative to the indicated LXG protein encoded by NCIB 3610 (y-axis). The competition outcome of the different wild isolates when cocultured with NCIB 3610 Δ*rapP* which has an equivalent doubling time is shown (data from Figure 2A). (D) Interaction outcomes with Δ*rapP* Δ*comA* (gfp+) (NRS7771) as challenger and the indicated strains (mTagBFP+). ****p < 0.0001, Significance determined by Welch’s t test.

Therefore, to investigate the impact of *ywqJ* in mediating Δ*rapP* Δ*comA* competitive fitness, we constructed a deletion of *ywqJ* in NCIB 3610 (mTagbfp+) (NRS8185). A dual isolate biofilm competition assay between Δ*rapP* Δ*comA* (gfp+) (NRS7771) and the *ywqJ* mutant displayed higher survival rates for Δ*rapP* Δ*comA* (gfp+) compared to a competition assay between Δ*rapP* Δ*comA* (gfp+) and the parental NCIB 3610 (gfp+) strain (Figure 5D, Figure S5C). Collectively, these results indicate that *ywqJ* contributes to intraspecies competition, but it is likely not to be the sole determinant of the interaction outcome as the two competing strains did not remain in equal prevalence after co-culture suggesting the involvement of other factors.

### Mutations within *comP* are detected in the genomes of *B. subtilis* isolates

Given the propensity of *comP* to acquire mutations during our evolution experiments, and the competitive advantage associated with such changes, we asked whether *comP* mutations were present in natural populations of *B. subtilis*. We analysed 370 high-quality genomes (See Supplemental File F2, Table S4), placed them within a phylogenetic framework (Figure 6A), and extracted *comQXPA* sequences with a focus on *comP*. As the *comQXP* region is highly polymorphic (Ansaldi and Dubnau, 2004), we screened specifically for (i) internal nonsense mutations causing premature truncation, (ii) frameshift-inducing indels, (iii) in-frame deletions within the conserved 3′ region of *comP*, and (iv) transposon insertions (28). Using these criteria, we identified 60 genomes carrying disrupted *comP* alleles, representing at least 16% of the dataset. This figure is likely an underestimate, as point mutations disrupting ComP function were not included. To validate this classification, we sourced seven strains from stock centres: four predicted to carry an intact *comP* allele (GCA_029537135, DSM 13109; GCA_037914925, DSM 10; GCA_024507895, BGSC 10A5; GCA_024508175, BGSC B98af) and three predicted to carry disrupted alleles (GCA_029536975, DSM 5611; GCA_029537155, DSM 1090; GCA_004101945, ATCC 11774/NCTC 8236). Short-read sequencing confirmed the predicted *comP* status in all cases, supporting the accuracy of our bioinformatic screen (See Supplemental File F2, Table S1).

**Figure 6:**
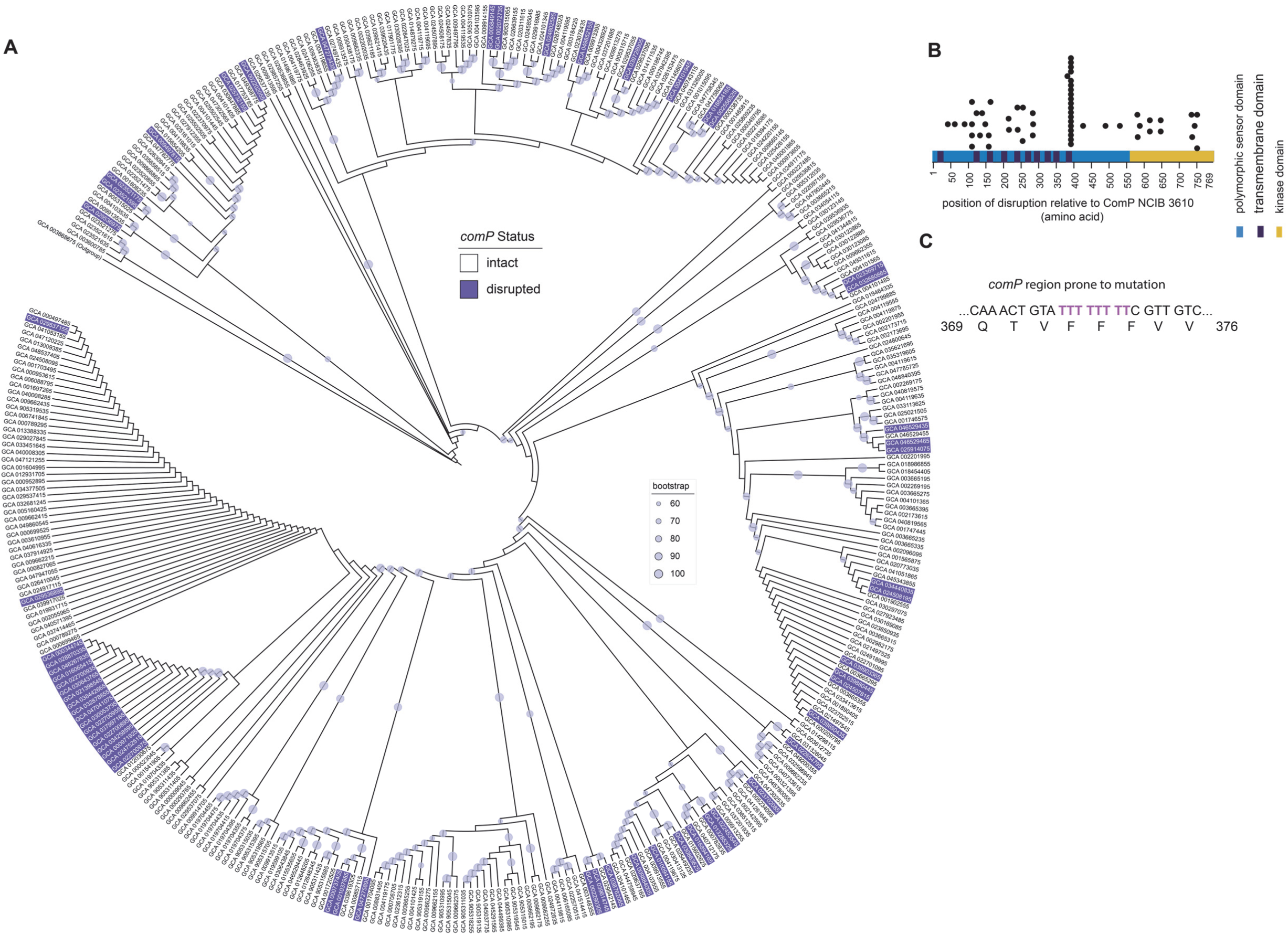
Distribution of mutations in the *B. subtilis comP* coding region. (A) 370 genomes of *B. subtilis* with *comP* status indicated as intact or disrupted. (B) The position of the *comP* disruption with respect to the ComP amino acid sequence is indicated with the calculated length of the variant protein produced (using NCIB 3610 ComP sequence as the reference) indicated by a black circle. (C) The string of eight thymidine found in wild type *comP* that form a mutation hotspot is indicated. This sequence is not found in all pherotypes of *comQXPA*.

Computational analysis of the 60 *comP* mutations showed that disruptive changes occurred throughout the coding sequence (Figure 6B) and were distributed broadly across the phylogeny. One notable cluster of closely related strains carried mutations within a poly-thymidine tract (eight consecutive T residues; Figure 6C), likely reflecting a mutational hotspot caused by slipped-strand mispairing during replication. These 18 strains were isolated from a wide range of environments between 2004 and 2023, including rhizosphere soil (wheat, cucumber), plant tissues (e.g., *Passiflora edulis*, tobacco leaves), aquatic and anthropogenic sources (river water, aquaculture ponds, sewage sludge, human faeces), and marine invertebrates (gills of *Teredo navalis*). Most isolates originated from multiple provinces in China (Guangxi, Henan, Jiangsu, Guangzhou, Shanxi, Changzhou, Lanzhou, Baoding, Anshun), with additional isolates from agricultural soil in Canada and from South Korea. Thus, this cluster of isolates reflects both ecological and geographic diversity while retaining phylogenetic similarity.

Finally, we assigned isolates to ComQXPA pherotypes based on ComQ sequence (Figure S6; (15)). Strains within the *comP*-mutant cluster all belonged to the NCIB 3610 ComQ clade, indicating a shared pherotype. Sequence divergence in the 5′ coding region of other pherotypes prevents the formation of an equivalent poly-thymidine tract, explaining the apparent restriction of this hotspot to the NCIB 3610 clade (see Supplemental File F4). Collectively, our analysis demonstrates that *comP* mutations are widespread across natural and anthropogenic environments, occur across phylogenetic groups, and can arise convergently within specific pherotypes.

## Discussion

### Quorum-sensing disruption as a strategy for competitive advantage

Intraspecies competition is a fundamental force shaping microbial community structure and evolution. Here we show that in *B. subtilis,* disruption of key quorum sensing pathways, either through natural variation or experimental evolution, confers a significant fitness advantage, even in structured biofilm communities where evolutionary pressure might be expected to favour communal traits. Specifically, we demonstrate that the RapP-PhrP module in strain NCIB 3610 and spontaneous mutations in the *comP* sensor kinase promote increased growth and competitive exclusion of otherwise closely related strains. However, growth is not the only determinant of the competition outcome as isolates with comparable growth rates still show antagonism. Antagonistic interactions were influenced by the profile of Type VII secretion system substrates.

In NCIB 3610, the RapP-PhrP^3610^ system is rendered unresponsive to its cognate signal due to a single amino acid substitution in RapP, leading to sustained inhibition of ComA (20). The N236T mutation in RapP effectively disables signal responsiveness, decoupling quorum sensing from cell density (20). This rewiring suppresses ComQXPA-dependent signalling, accelerates growth (20, 22) and enhances competitive ability in biofilm environments. It also renders NCIB 3610’s behaviour atypical relative to true wild strains, particularly in social traits like secondary metabolite production, swarming, and biofilm formation (21)(Omer Bendori et al., 2015; Parashar et al., 2013). We therefore stress the conclusion that NCIB 3610, while genetically undomesticated in some respects (it is not an auxotroph), should be considered functionally domesticated in terms of its social regulatory behaviour (23).

### Experimental evolution and parallels across bacterial systems

Experimental evolution of strains typically outcompeted by NCIB 3610 repeatedly yielded mutations in *comP* that mimic the effects of RapP^3610^; that is, reduced ComA activity, increased growth, and improved fitness during co-culture with NCIB 3610.

These findings underscore a central theme: the suppression of communal signalling pathways can favour individual competitiveness. This trade-off between quorum-sensing–regulated cooperation and rapid growth is increasingly recognised across bacterial systems. In *Pseudomonas aeruginosa*, for example, *lasR* mutants that disrupt quorum sensing frequently emerge during chronic infections, displaying enhanced growth while losing virulence factor production and public goods sharing (29, 30). Additionally, in *Streptomyces* the inactivation of costly antibiotic biosynthesis pathways improves growth under nutrient-limiting conditions (31).

Spontaneous quorum sensing-defective mutants have been observed in diverse bacteria during laboratory culture, including *Burkholderia glumae* (32) and *P. aeruginosa* (33). Consistent with these examples, our findings show that disruption of the *comQXPA* or *rapP-phrP* systems can offer a competitive edge in structured microbial communities. Importantly, such quorum sensing disruptions are not restricted to experimental evolution. Our analysis of 370 *B. subtilis* genomes shows that disruptive *comP* mutations are not rare but present in at least 16% of wild isolates, particularly in those sharing the NCIB 3610 ComQXPA pherotype. This suggests a recurrent and naturally selected strategy whereby quorum sensing is attenuated to reduce the cost of social engagement under competitive pressure.

Our analysis further explores the complex regulatory interplay between the Rap–Phr and ComQXPA systems (Figure 1A). The competitive advantage conferred by RapP^3610^ is only partially dependent on ComA, as evidenced by the intermediate phenotype of the *rapP comA* double mutant. RapP also influences Spo0A activity by dephosphorylating Spo0F, thereby altering matrix production and sporulation (4).

Moreover, during the evolution experiments we isolated a derivative of NRS6145 with a deletion of *rhgR1* (Figure 3), which encodes a repressor of several Rap phosphatases including RapD, RapG, and RapH (34). Loss of *rhgR1* would be expected to derepress these Rap proteins, enabling them to dephosphorylate their targets and perhaps further modulate ComA and Spo0A activity. Thus, in the NRS6145-L6P4 *rhgR1* containing mutant, multiple Rap-mediated inputs may be simultaneously deregulated, expanding the regulatory pathways that shape competitive outcomes. As Rap-Phr systems integrate environmental cues such as nutrient availability and extracellular peptide signals (20, 35), perturbations at this level are likely to propagate broadly.

### Beyond growth: the role of antagonistic effectors

While growth rate is a strong predictor of competitive fitness, our findings make clear that it is not the sole determinant. Strains with equivalent doubling times exhibited contrasting outcomes depending on their regulatory configuration. For instance, NCIB 3610 and its *rapP comA* double mutant grew at similar rates in planktonic culture, but only the parental strain dominated in biofilms. Using RNA sequencing analysis, we were able to narrow one of the contributors of the competition outcome to differential expression of the Type VII secretion substrate YwqJ. Toxin-mediated interference, such as through LXG-domain proteins, can confer dominance even at slower growth rates (10, 13), and differential expression on deletion of *rapP* and *comA* is consistent with the knowledge that the production of antibiotics or matrix components, often ComA-regulated (36, 37), can tip the balance depending on ecological context.

### Context-dependent fitness of quorum-sensing mutants

Crucially, the benefit of disabling ComA-based quorum sensing appears to be context-dependent. In our dual-*B. subtilis* strain model, where all competitors belong to the same species and are relatively resistant to each other’s secondary metabolites, the energetic cost of producing ComA-regulated traits likely outweighs their utility. In contrast, it has recently been shown that *B. subtilis* invests more heavily in secondary metabolites when confronted with phylogenetically distant species (26), precisely when such compounds are more likely to be effective.

Conversely, *B. subtilis* evolved under fungal competition (*Setophoma terrestris*) acquired mutations in *comQXPA* that led to increased production of antifungal volatiles, enhancing inter-kingdom fitness (38). These findings underscore that while quorum sensing repression is beneficial in intraspecies competition, it may be detrimental in broader ecological settings where toxin production is essential for survival.

Taken together, our study reveals that modulation of quorum sensing via genetic disruption or natural variation represents a powerful mechanism for navigating social conflict in microbial populations. It highlights the evolutionary plasticity of regulatory networks and suggests that the ecological value of cooperation is highly contingent on community composition and environmental structure. A systems-level understanding that integrates growth dynamics, regulatory complexity, spatial ecology, and inter-strain antagonism will be essential to predict and manipulate microbial community outcomes.

## Materials and Methods

### Growth Media and additives

For routine growth of *Bacillus subtilis,* lysogeny broth (LB) liquid media was made using the following recipe: 1% (w/v) Bacto-peptone, 1% (w/v) NaCl, 0.5% (w/v) yeast extract. For solid plates, LB broth was supplemented with 1.5% (w/v) agar. When necessary, LB media cultures and plates were supplemented with antibiotics which were used at the following concentrations for *B. subtilis*: 10 μg/ml kanamycin, 100 μg/ml spectinomycin, 5 μg/ml chloramphenicol and 0.5 μg/ml erythromycin combined with 12.5 μg/ml lincomycin for “MLS”. Ampicillin was used at 100 μg/ml for work with *E. coli*.

### Plasmid and Strain Construction

The strain used for storing of plasmids for cloning was *Escherichia coli* strain MC1061 [F’ *lacIQ lacZM15 Tn*10. For making mutations in the NCIB 3610 background, as this strain is not genetically competent, plasmids were first transformed into the laboratory strain 168 using standard protocols (Supplemental Methods).

### Markerless in-frame deletion strain construction

For construction of strains with in-frame deletions without the insertion of antibiotic resistance cassettes into their genomes, pMiniMAD-based plasmids (39) were used as detailed for the parental pMAD plasmid (40) (see Supplemental File F2, Table S6, Supplemental Methods).

### Setting up single strain colony biofilms

*B. subtilis* isolates were streaked out on LB agar plates and incubated at 37°C overnight. The following day single colonies were grown in 3 ml of LB broth at 37°C with agitation. The cultures were grown to an OD600 of 1 and 10 µl of the cultures were spotted onto MSgg media plates. The plates were incubated at 30°C for 48 h before imaging. Biofilm imaging was performed using a Leica MZ16 FA stereoscope, LAS version 2.7.1. The imaging data were imported to OMERO (41).

### Biofilm co-culture assays and surface area calculation

For dual isolate biofilm assays, cultures of the individual strains to be used were set up in 5 ml of LB and incubated at 37°C with agitation overnight. The following morning, day cultures were set up by inoculating 3 ml of LB with 200 µl of the overnight cultures. The day cultures were incubated at 37°C with agitation. The growth of the cultures was monitored, and all cultures were normalised to an OD600 of 1. After normalisation, cultures were mixed at a 1:1 ratio as required. 5 µl drops of the culture mixtures were spotted onto MSgg agar plates and 5 µl of the individual normalised cultures were included as controls. The plates were incubated at 30°C and imaged after 24, 48 and 72 hours as required. Fluorescence imaging was performed using a Leica fluorescence stereoscope (M205FCA) with a 0.5 × 0.2 NA objective. Imaging files were imported to OMERO (41). To quantify the biofilm footprint, we used OMERO.Insight v.5.8.1 (Allan et al., 2012). Each biofilm was highlighted as a region of interest, and the area occupied was calculated. Relative strain densities of GFP and mTagBFP-expressing cells in mixed biofilm assays were determined by analysing fluorescence image data (24). ImageJ/Fiji (42) was used to run the macro as previously described (24).

### Confocal Analysis of Colony Biofilms

Confocal imaging was done using methodology similar to that described previously (43). See Supplemental methods.

### Recovery of isolates from dual isolate colony biofilms

A colony biofilm was removed from the agar plate with a sterile plastic loop and placed in 500 µL of phosphate-buffered saline (PBS): 137 mM NaCl, 2.7 mM KCl, 8 mM Na2HPO4, and 2 mM KH2PO4. The biofilm was disrupted by passage through a 1 mL syringe with a 23G needle 6 times, a process repeated with a 27G needle.

Before plating, the disrupted biofilm was serially diluted in PBS and 100 µL was spread onto LB agar plates containing the antibiotic to which the desired strain was resistant. Colonies recovered were counted after overnight incubation at 37°C and enumerated where required.

### Isolation of cells from dual isolate colony biofilms for flow cytometry

Mixed isolate colony biofilms were set up by growing the cultures to an OD600 of 1 and mixing the strains of interest at a 1:1 ratio as described above. After 48h, a colony biofilm was removed from the agar plate with a sterile plastic loop and placed in 500 µL of 4% formaldehyde before processing (Supplemental Methods).

### Experimental evolution

Strains NRS6936, NRS6951, and NRS7201 (NRS6105, NRS6145 and NRS6202 respectively adapted to express mTagBFP and containing spectinomycin resistant) and NRS1473 (NCIB 3610 expressing GFP, kanamycin resistant), were used for the experimental evolution detailed in the Supplemental Methods.

### Genomic DNA extraction and sequencing

Genomic DNA was extracted using the QIAprep® Spin Miniprep Kit (cat. Nos. 27104 and 27106) and following the “Quick-Start Protocol”. DNA concentration was measured using a Qubit 4 Fluorometer or a NanoDrop™ One/One^C^ Microvolume UV-Vis Spectrophotometer. Whole Genome Sequencing was performed by GENEWIZ Germany GmbH (Part of Azenta Life Sciences), MicrobesNG (UK) or SeqCentre (USA) as indicated in the strain table (See Supplemental File F2, Table S1). See Supplemental Methods for more information.

### Bacillus subtilis phylogenetic tree

Sequence selection and preparation for phylogenetic analysis was conducted using custom Python code co-ordinated through a Snakemake workflow (44). A database of ‘complete’ genome *B. subtilis* sequences was constructed from the European Nucleotide Archive (with the genome represented in <5 contigs, to allow for plasmid sequences) which had been assigned an NCBI taxonomy ID of 1423, including child taxa, resulting in a set of 478 genomes (Data downloaded on 6^th^ June 2025) (see Supplemental File F2, Table S4). Genome sequences were reannotated with Bakta 1.11.0 (45) (database version 5.1) to ensure consistent annotation. Genome completeness was assessed using Busco 5.8.3 (46) using the ‘bacillales_odb10’ lineage, and the database filtered to remove genomes with a Busco completeness of <98%. Taxonomic classification was carried out using GTDB-TK 2.4.1 (GTDB reference data version r226) (47) and misclassified sequences removed from the database, retaining only those classified as *B. subtilis*. The final database contained 370 genome sequences.

### Analysis of ComP sequences in *Bacillus subtilis* genomes sequences

Operon sequences from the start of ComQ to the end of ComA were extracted from the annotated genome sequences and analysed (Supplemental Methods).

### RapP and LXG protein sequence analysis

Proteins in the pfam family PF04740 LXG domain of WXG superfamily (48) were identified in each strain using HMMER (49).

### Measurement of Growth Rate

To calculate the doubling time of cultures, strains were streaked out and incubated for 16 hours at 30°C. Liquid overnight cultures were established in 5 ml of LB and incubated at 37°C with agitation at 250 rpm for 16 hours. Day cultures were established by dilution (200 μl into 5 ml) and grown at 37°C with shaking for 3 hours and 10 mins. The tubes were centrifuged, and the cell pellets were resuspended in 1 ml of MSgg. The OD600 was measured, and the culture normalised to an OD600 of 1 using MSgg. Cultures for growth measurement were set up to an initial OD600 of 0.01 using 25 ml of pre-warmed MSgg and incubated at 30°C with 250 rpm agitation.

OD600 values were recorded every 30-60 minutes the doubling time calculated using the exponential growth phase measurements.

### Structural modelling of ComP

The protein sequences of ComP from strain NRS6145 (wild type, WT) and the mutant variant carrying the E510K substitution were submitted to the AlphaFold2 server (25) (see Supplemental File F2, Table S8, Supplemental Methoda)

### RNA isolation and RNAseq analysis

RNAseq analysis was conducted on strains NRS6942 and NRS7771, grown for 20 h at 30°C on MSgg agar plates. Five biological replicates were used, using 4 technical replicates pooled from different plates. The RNA isolation protocol was adapted (50) using a Qiagen RNeasy RNA Isolation kit (See Supplemental Methods). Library construction and RNA sequencing was performed by Edinburgh Genetics using Stranded Total RNA Prep with Ribo-Zero Plus Microbiome kit. RNA was sequenced using Illumina Paired Ends 100bp, with >=12M PE reads per sample. The data processing is detailed in the supplemental methods.

### Mathematical model of dual-isolate biofilms

We used a mathematical model to test hypotheses on the impact of an isolate’s doubling time on its competitiveness during biofilm growth. Adapted from an existing mathematical model (24), the model describes the spatio-temporal dynamics of two isolates in a single growing biofilm (Supplemental Methods).

## Data Analysis

Graphs were constructed using GraphPad Prism 9 or R/ggplot2. Statistical analysis methods are detailed in the legends.

## Acknowledgements

Work was funded by the UKRI Biotechnology and Biological Science Research Council (BBSRC) [BB/P001335/1, BB/R012415/1, BB/Z516600/1, BB/X002950/1].

MK and AB were supported by a UKRI Biotechnology and Biological Sciences Research Council studentship [BB/M010996/1, BB/T00875X/1]. AW is supported by a Wellcome PhD fellowship [218520/Z/19/Z]. HCD is supported by a Wellcome Career Development Award 225350/Z/22/Z.

We thank Dr. Daniel Neill, Prof. Susanne Gebhard and Prof Marie Elliot for review of the draft manuscript. We thank members of the Dundee Imaging Facility and Flow Cytometry Facility (Dr. Clarke) for assistance with experiments. We thank Prof Dan Kearns for sharing plasmid pMP68.

## Declarations

Some of the work presented in the manuscript has been presented in the PhD thesis of Margarita Kalamara published by the University of Dundee.

ChatGPT was used to edit an R script used to generate Figure 5C and for help in refining scripts used to assemble genomes.

For the purpose of open access, the author has applied a CC BY public copyright licence to any Author Accepted Manuscript version arising from this submission.

## Data Availability Statement

Code used to analyse the data and generate figures are available at https://doi.org/10.5281/zenodo.17463645 (51). Experimental datasets have been deposited at https://doi.org/10.5281/zenodo.17463729 (52) and https://www.ebi.ac.uk/biostudies/studies/S-BSST2262. Interactive phylogenetic trees are available through the Interactive Tree of Life: https://itol.embl.de/shared/2mG3crJad7HbC

## REFERENCES

1. L. Philippot, B. S. Griffiths, S. Langenheder, Microbial Community Resilience across Ecosystems and Multiple Disturbances. Microbiol Mol Biol Rev 85 (2021).

2. S. Liu et al., Decoding bacterial communication: Intracellular signal transduction, quorum sensing, and cross-kingdom interactions. Microbiological research 292, 127995 (2025).

3. W. Tan, H. Nian, L. P. Tran, J. Jin, T. Lian, Small peptides: novel targets for modulating plant-rhizosphere microbe interactions. Trends Microbiol 32, 1072–1083 (2024).

4. S. Arnaouteli, N. C. Bamford, N. R. Stanley-Wall, A. T. Kovacs, Bacillus subtilis biofilm formation and social interactions. Nat Rev Microbiol 19, 600–614 (2021).

5. A. Dragos et al., Division of Labor during Biofilm Matrix Production. Curr Biol 28, 1903–1913 e1905 (2018).

6. K. Kobayashi, Y. Ikemoto, Biofilm-associated toxin and extracellular protease cooperatively suppress competitors in Bacillus subtilis biofilms. PLoS Genet 15, e1008232 (2019).

7. M. J. Powers, E. Sanabria-Valentin, A. A. Bowers, E. A. Shank, Inhibition of Cell Differentiation in Bacillus subtilis by Pseudomonas protegens. J Bacteriol 197, 2129–2138 (2015).

8. A. A. Schoenborn et al., Defining the Expression, Production, and Signaling Roles of Specialized Metabolites during Bacillus subtilis Differentiation. J Bacteriol 203, e0033721 (2021).

9. S. M. Yannarell, D. Velickovic, C. R. Anderton, E. A. Shank, Direct Visualization of Chemical Cues and Cellular Phenotypes throughout Bacillus subtilis Biofilms. mSystems 6, e0103821 (2021).

10. N. A. Lyons, R. Kolter, A single mutation in rapP induces cheating to prevent cheating in Bacillus subtilis by minimizing public good production. Commun Biol 1, 133 (2018).

11. N. A. Lyons, B. Kraigher, P. Stefanic, I. Mandic-Mulec, R. Kolter, A Combinatorial Kin Discrimination System in Bacillus subtilis. Curr Biol 26, 733–742 (2016).

12. P. Stefanic et al., Kin discrimination promotes horizontal gene transfer between unrelated strains in Bacillus subtilis. Nat Commun 12, 3457 (2021).

13. K. Kobayashi, Diverse LXG toxin and antitoxin systems specifically mediate intraspecies competition in Bacillus subtilis biofilms. PLoS Genet 17, e1009682 (2021).

14. M. Kalamara, J. Abbott, T. Sukhodub, C. MacPhee, N. R. Stanley-Wall, The putative role of the epipeptide EpeX in Bacillus subtilis intra-species competition. Microbiology (Reading) 169 (2023).

15. P. Stefanic, I. Mandic-Mulec, Social interactions and distribution of Bacillus subtilis pherotypes at microscale. J Bacteriol 191, 1756–1764 (2009).

16. P. Stefanic et al., The quorum sensing diversity within and between ecotypes of Bacillus subtilis. Environ Microbiol 14, 1378–1389 (2012).

17. M. Perego, J. A. Hoch, Cell-cell communication regulates the effects of protein aspartate phosphatases on the phosphorelay controlling development in Bacillus subtilis. Proc Natl Acad Sci U S A 93, 1549–1553 (1996).

18. B. A. Lazazzera, I. G. Kurtser, R. S. McQuade, A. D. Grossman, An autoregulatory circuit affecting peptide signaling in Bacillus subtilis. J Bacteriol 181, 5193–5200 (1999).

19. R. Gallegos-Monterrosa, A. T. Kovacs, Phenotypic plasticity: The role of a phosphatase family Rap in the genetic regulation of Bacilli. Mol Microbiol 120, 20–31 (2023).

20. S. Omer Bendori, S. Pollak, D. Hizi, A. Eldar, The RapP-PhrP quorum-sensing system of Bacillus subtilis strain NCIB3610 affects biofilm formation through multiple targets, due to an atypical signal-insensitive allele of RapP. J Bacteriol 197, 592–602 (2015).

21. V. Parashar, M. A. Konkol, D. B. Kearns, M. B. Neiditch, A plasmid-encoded phosphatase regulates Bacillus subtilis biofilm architecture, sporulation, and genetic competence. Journal of Bacteriology 195, 2437–2448 (2013).

22. M. Zhu et al., Plasmid-encoded phosphatase RapP enhances cell growth in non-domesticated Bacillus subtilis strains. Nat Commun 15, 9567 (2024).

23. S. Pollak, S. Omer Bendori, A. Eldar, A complex path for domestication of B. subtilis sociality. Curr Genet 61, 493–496 (2015).

24. L. Eigentler et al., Founder cell configuration drives competitive outcome within colony biofilms. ISME J 10.1038/s41396-022-01198-8 (2022).

25. J. Abramson et al., Accurate structure prediction of biomolecular interactions with AlphaFold 3. Nature 630, 493–500 (2024).

26. H. Maan, M. Itkin, S. Malitsky, J. Friedman, I. Kolodkin-Gal, Resolving the conflict between antibiotic production and rapid growth by recognition of peptidoglycan of susceptible competitors. Nature Communications 13 (2022).

27. T. Pedreira, C. Elfmann, J. Stulke, The current state of SubtiWiki, the database for the model organism Bacillus subtilis. Nucleic Acids Res 50, D875–D882 (2022).

28. T. Nagai, L. S. Tran, Y. Inatsu, Y. Itoh, A new IS4 family insertion sequence, IS4Bsu1, responsible for genetic instability of poly-gamma-glutamic acid production in Bacillus subtilis. J Bacteriol 182, 2387–2392 (2000).

29. S. P. Diggle, A. S. Griffin, G. S. Campbell, S. A. West, Cooperation and conflict in quorum-sensing bacterial populations. Nature 450, 411–414 (2007).

30. J. B. Feltner et al., LasR Variant Cystic Fibrosis Isolates Reveal an Adaptable Quorum-Sensing Hierarchy in Pseudomonas aeruginosa. mBio 7 (2016).

31. S. Westhoff, A. M. Kloosterman, S. F. A. van Hoesel, G. P. van Wezel, D. E. Rozen, Competition Sensing Changes Antibiotic Production in Streptomyces. mBio 12 (2021).

32. G. Gnanasekaran, J. Y. Lim, I. Hwang, Disappearance of Quorum Sensing in Burkholderia glumae During Experimental Evolution. Microb Ecol 79, 947–959 (2020).

33. S. Robitaille, M. C. Groleau, E. Deziel, Swarming motility growth favours the emergence of a subpopulation of Pseudomonas aeruginosa quorum-sensing mutants. Environ Microbiol 22, 2892–2906 (2020).

34. K. Hayashi, T. Kensuke, K. Kobayashi, N. Ogasawara, M. Ogura, Bacillus subtilis RghR (YvaN) represses rapG and rapH, which encode inhibitors of expression of the srfA operon. Mol Microbiol 59, 1714–1729 (2006).

35. S. Pollak et al., Facultative cheating supports the coexistence of diverse quorum-sensing alleles. Proc Natl Acad Sci U S A 113, 2152–2157 (2016).

36. M. Ogura, H. Yamaguchi, K. Yoshida, Y. Fujita, T. Tanaka, DNA microarray analysis of Bacillus subtilis DegU, ComA and PhoP regulons: an approach to comprehensive analysis of B.subtilis two-component regulatory systems. Nucleic Acids Res 29, 3804–3813 (2001).

37. J. M. Auchtung, C. A. Lee, A. D. Grossman, Modulation of the ComA-dependent quorum response in Bacillus subtilis by multiple Rap proteins and Phr peptides. J Bacteriol 188, 5273–5285 (2006).

38. A. G. Albarracin Orio et al., Fungal-bacterial interaction selects for quorum sensing mutants with increased production of natural antifungal compounds. Commun Biol 3, 670 (2020).

39. K. J. Wozniak, L. A. Simmons, Genome Editing Methods for Bacillus subtilis. Methods Mol Biol 2479, 159–174 (2022).

40. M. Arnaud, A. Chastanet, M. Debarbouille, New vector for efficient allelic replacement in naturally nontransformable, low-GC-content, gram-positive bacteria. Appl Environ Microbiol 70, 6887–6891 (2004).

41. C. Allan et al., OMERO: flexible, model-driven data management for experimental biology. Nat Methods 9, 245–253 (2012).

42. J. Schindelin et al., Fiji: an open-source platform for biological-image analysis. Nat Methods 9, 676–682 (2012).

43. M. Porter, F. A. Davidson, C. E. MacPhee, N. R. Stanley-Wall, Systematic microscopical analysis reveals obligate synergy between extracellular matrix components during Bacillus subtilis colony biofilm development. Biofilm 4 (2022).

44. F. Molder et al., Sustainable data analysis with Snakemake. F1000Res 10, 33 (2021).

45. O. Schwengers et al., Bakta: rapid and standardized annotation of bacterial genomes via alignment-free sequence identification. Microb Genom 7 (2021).

46. M. Manni, M. R. Berkeley, M. Seppey, E. M. Zdobnov, BUSCO: Assessing Genomic Data Quality and Beyond. Curr Protoc 1, e323 (2021).

47. P. A. Chaumeil, A. J. Mussig, P. Hugenholtz, D. H. Parks, GTDB-Tk v2: memory friendly classification with the genome taxonomy database. Bioinformatics 38, 5315–5316 (2022).

48. T. Paysan-Lafosse et al., InterPro in 2022. Nucleic Acids Res 51, D418–D427 (2023).

49. R. D. Finn, J. Clements, S. R. Eddy, HMMER web server: interactive sequence similarity searching. Nucleic Acids Res 39, W29–37 (2011).

50. M. Futo et al., Embryo-Like Features in Developing Bacillus subtilis Biofilms. Mol Biol Evol 38, 31–47 (2021).

51. N. Stanley-Wall, J. C. Abbott, Regulatory-Rewiring-Drives-Intraspecies-Competition-in-Bacillus-subtilis - Software. Zendod.

52. N. Stanley-Wall, Regulatory Rewiring Drives Intraspecies Competition in Bacillus subtilis. Zenodo.

