## Supplemental data for "Regulatory Rewiring Drives Intraspecies Competition in *Bacillus subtilis*"

**Supplemental Methods**

**Supplemental Figures**

**Supplemental References**

**Supplementary File List**

**File F1. Breseq Analysis**

**File F2. Supplementary Tables S1-S8**.

- Table S1: Strains used in this study
- Table S2: Functional terms/pathways derived from GO-terms and KEGG pathways
- Table S3: List of 89 toxins and antibiotics. Gene symbols provided were matched to the SubtiWiki gene symbols, mapped to NCBI annotations.
- Table S4: Genomic sequences used in this study
- Table S5: Primers used in this study.
- Table S6: Plasmids used in this study.
- Table S7: Sequences of constructs synthesized.
- Table S8. Protein sequences used in this study.

**File F3. Jalview com alignments.**

**File F4. Jalview ywqJ alignments.**

| Term | Definition | Initials |
|---|---|---|
| Conceptualization | Ideas; formulation or evolution of overarching research goals and aims | NSW, MK |
| Methodology | Development or design of methodology; creation of models | NSW, JCA, LE, MP, MK, AB |
| Software | Programming, software development; designing computer programs; implementation of the computer code and supporting algorithms; testing of existing code components | JCA, MG, LE, NSW |
| Validation | Verification, whether as a part of the activity or separate, of the overall replication/ reproducibility of results/experiments and other research outputs | NSW, MK |
| Formal analysis | Application of statistical, mathematical, computational, or other formal techniques to analyze or synthesize study data | NSW, MG, JCA, LE, MK, AB, JC |
| Investigation | Conducting a research and investigation process, specifically performing the experiments, or data/evidence collection | NSW, JG, DS, MK, AB, JC |
| Resources | Provision of study materials, reagents, materials, patients, laboratory samples, animals, instrumentation, computing resources, or other analysis tools | NSW, JCA |
| Data Curation | Management activities to annotate (produce metadata), scrub data and maintain research data (including software code, where it is necessary for interpreting the data itself) for initial use and later reuse | NSW, JG, JCA, MG, LE, AB |
| Writing - Original Draft | Preparation, creation and/or presentation of the published work, specifically writing the initial draft (including substantive translation) | NSW, JG, JCA, MK, AB |
| Writing - Review & Editing | Preparation, creation and/or presentation of the published work by those from the | NSW, CEM, JG, JCA, MG, LE, MP, MK, AB, JC |

| Term | Definition | Initials |
|---|---|---|
| | original research group, specifically critical review, commentary or revision – including pre-or postpublication stages | |
| Visualization | Preparation, creation and/or presentation of the published work, specifically visualization/ data presentation | NSW, JG, JCA, MG, MK, AB |
| Supervision | Oversight and leadership responsibility for the research activity planning and execution, including mentorship external to the core team | NSW, CEM |
| Project administration | Management and coordination responsibility for the research activity planning and execution | NSW |
| Funding acquisition | Acquisition of the financial support for the project leading to this publication | NSW, CEM |

51

### Supplemental Methods

#### *MSgg biofilm-inducing media*

MSgg (Minimal Salts glycerol glutamate) medium was made by making a base medium, consisting of 5 mM potassium phosphate, 100 mM MOPS at pH=7.0, supplemented with 1.5% (w/v) agar. The media base was autoclaved and cooled to 55°C. The base medium was supplemented with 2 mM $MgCl_2$, 700 µM $CaCl_2$, 50 µM $FeCl_3$, 50 µM $MnCl_2$, 1 µM $ZnCl_2$, 2 µM thiamine, 0.5% (v/v) glycerol and 0.5% (w/v) glutamic acid. A volume of 23 ml of molten MSgg was added to each 9 cm diameter petri dish and the plates were solidified at room temperature. The solidified plates were dried for 1 hour under a laminar flow cabinet to dry before use.

#### *TY media for phage preparation and transduction*

For phage propagation, LB was supplemented with 10 mM $MgSO_4$ and 1 µM $MnSO_4$ to make TY media. TY top agar was made by the addition of agar at a 0.7% (w/v) concentration and agar plates were made by the addition of 1.5% (w/v) agar. For transduction of phage into cells, to prevent cell lysis, TY agar plates were supplemented with 10 mM sodium citrate.

#### *Modified Competence media (MC) for genetic modification of B. subtilis*

Modified media (MC) was made at a 10 x concentration as previously described (1) with the following adaptations. The solution was made by mixing 10.7 g $K_2HPO_4$, 5.2 g $KH_2PO_4$, 20 g dextrose, 0.88 g sodium citrate dehydrate, 2.2 g L-glutamic acid monopotassium salt and 1 g casein enzymatic hydrolysate per 100 ml of solution, dissolved in milli-Q water. The solution was sterilised by passage through a 0.2 µm pore size membrane filter. Before use, the concentrated medium was diluted to 1x concentration in milli-Q water and supplemented with 30 µl of 1 M $MgSO_4$ and 17.5 µl of 500 mM $FeCl_3$ per 10 ml of media.

#### *Plasmid and Strain Construction*

The strain used for storing of plasmids for cloning was *Escherichia coli* strain MC1061 [F' *lacIQ lacZM15 Tn*10. For making mutations in the NCIB 3610 background, as this strain is not genetically competent, plasmids were first transformed into the laboratory strain 168 using standard protocols. The modified region was subsequently inserted and integrated into the NCIB 3610 genome via SPP1 phage transduction. For genetically competent soil isolates of *B. subtilis*, the plasmids were transformed directly into the isolate of interest as previously described with the adaptations described. Plasmids were either constructed using standard restriction digest methods using the primers (see Supplemental File F2, Table S5) and backbone plasmids (see Supplemental File F2, Table S6) detailed or were synthesised (see Supplemental File F2, Table S7). Growth media conditions are detailed in the Supplemental Methods.

#### *Isolation of cells from dual isolate colony biofilms for flow cytometry*

Mixed isolate colony biofilms were set up by growing the cultures to an $OD_{600}$ of 1 and mixing the strains of interest at a 1:1 ratio as described above. After 48h, a colony biofilm was removed from the agar plate with a sterile plastic loop and placed in 500 µL of 4% formaldehyde before processing. The biofilm was disrupted by passage through a 1 µL syringe with a 23G needle 6 times and incubated at room temperature for 7 minutes. The cells were collected by centrifugation for 1 minute at 10,000 g and washed in 1 ml 1xPBS (137 mM NaCl, 2.7 mM KCl, 8 mM $Na_2HPO_4$, and 2 mM $KH_2PO_4$). The cells were harvested and suspended in 500 µL of GTA buffer (10 mM EDTA (pH=8), 20mM Tris-HCl (pH=8), 50mM glucose). Before analysis by flow cytometry, the samples were subjected to mild sonication using 5 cycles of 1 sec pulse and 3 seconds rest on ice at 30% power. 1 µl of the cells released from the biofilm was added to 200 µl of 4% (w/v) BSA in PBS that had been filtered twice and placed in a 96-well plate. Samples were analysed on an ID7000 spectral analyser (Sony Biosciences). Forward scatter (FSC) and side scatter (SSC) were detected from the 488nm laser with threshold set on SSC. GFP fluorescence was quantified using 488 nm excitation and the ‘virtual filter’ Channel 4-Channel 6 (493.9-530.3 nm).

#### *Markerless in-frame deletion strain construction*

For NCIB 3610 derived strains, the plasmid of interest was introduced into *B. subtilis* strain 168. This was done by mixing 100 µl of thawed competent 168 cells with 100 µl of transformation buffer and approximately 1.2 µg of plasmid DNA and incubated at 30$^{o}$C for 1 hour. The sample was plated on LB plates containing MLS for selection and incubated at 30°C for 48 hours. Multiple colonies were pooled into 5 ml of TY both containing MLS and incubated at 30°C with agitation overnight. This culture was used to make and isolate phage containing the plasmid as described above. The phage was transduced into NCIB 3610 as described above but the incubation step following the addition of phage to the TY culture was conducted at 30°C for 1 hour. After plating, on LB containing 10 mM sodium citrate and MLS antibiotics for selection was incubated at 30°C for 48 hours. Multiple transduced colonies were picked from the plates and inoculated into the same 5 ml LB culture containing MLS. The culture was grown at 37°C overnight with shaking, which is a non-permissive temperature for the replication of the plasmid that selects for integration of the plasmid into the chromosome due to the presence of the antibiotic in the media. The following day, the culture was serially diluted and plated onto LB plates containing MLS to select for cells that had undergone as successful integration of the plasmid into the genome. The plates were incubated at 37°C overnight. Next, multiple colonies were selected and inoculated into 5 ml of LB broth with no antibiotics. The culture was growing without shaking at 30°C overnight and then for 4 hours at 30°C with shaking. The culture was serially diluted and plated onto LB plates and incubated at 37°C overnight. Approximately 40 colonies were selected and replica-patched on LB plates and LB plates containing MLS. The patch plates were incubated at 37°C overnight and colonies that were sensitive to MLS were screened by colony PCR. For clones that appeared to have acquired the mutation of interest, the PCR product was sequenced to confirm the presence of the in-frame deletion on the chromosome (see Supplemental File F2, Table S5). For construction of in-frame deletions in the NRS6105 background, a genetically competent wild type isolate, the pMiniMAD- based plasmid (see Supplemental File F2, Table S6) was transformed into the isolate using the MC media transformation protocol but incubating the cells at 30°C for 3 hours following the addition of the plasmid. The transformed cells were plated onto LB plates containing MLS for selection and the plates were incubated at 30°C for 48 hours. Multiple colonies from the plates were picked and inoculated into 5 ml of LB containing MLS and the culture was incubated at 37°C overnight to induce integration of the plasmid into the chromosome. The rest of the process followed was the same as for NCIB 3610.

*Genomic DNA extraction and sequencing*

Illumina sequencing libraries were prepared by SeqCentre using the tagmentation-based and PCR-based Illumina DNA Prep kit and custom IDT 10bp unique dual indices (UDI) with a target insert size of 280 bp. No additional DNA fragmentation or size selection steps were performed. Illumina sequencing was performed on an Illumina NovaSeq X Plus sequencer in one or more multiplexed shared-flow-cell runs, producing 2x151bp paired-end reads. Demultiplexing, quality control and adapter trimming was performed with bcl-convert1 (v4.2.4).

For Illumina sequencing by MicrobesNG, genomic DNA libraries were prepared using the Nextera XT Library Prep Kit (Illumina, San Diego, USA) following the manufacturer's protocol with the following modifications: input DNA was increased 2-fold, and PCR elongation time was increased to 45 s. DNA quantification and library preparation were carried out on a Hamilton Microlab STAR automated liquid handling system (Hamilton Bonaduz AG, Switzerland). Pooled libraries were quantified using the Kapa Biosystems Library Quantification Kit for Illumina. Libraries were sequenced using Illumina sequencers (HiSeq/NovaSeq) using a 250 bp paired end protocol.

For sequencing by Genewiz, genomic DNA samples were quantified using Qubit 4.0 Fluorometer (Life Technologies, Carlsbad, CA, USA) and DNA integrity was checked using Agilent TapeStation 4200 (Agilent Technologies, Palo Alto, CA, USA). NEBNext Ultra II DNA Library Preparation kit for was used for DNA library preparation following the manufacturer's recommendations (NEB, Ipswich, MA, USA). Briefly, genomic DNA was fragmented by acoustic shearing with a Covaris instrument. Fragmented DNA was end repaired and adenylated. Adapters were ligated after adenylation of the 3'ends followed by enrichment by limited cycle PCR. Adapter-ligated DNA libraries were cleaned up and validated using Agilent TapeStation and quantified using Qubit 4.0 Fluorometer. The libraries have been multiplexed on a flowcell and loaded on the Illumina NovaSeq 6000 instrument according to the manufacturer's instructions. The samples were sequenced using a 2x150 paired-end (PE) configuration. Image analysis and base calling was conducted by the NovaSeq Control Software on the NovaSeq instrument. Raw sequence data (.bcl files) generated from Illumina NovaSeq was converted into fastq files and de-multiplexed using Illumina bcl2fastq program version 2.20. One mismatch was allowed for index sequence identification.

Reads were adapter trimmed using Trimmomatic 0.30 with a sliding window quality cutoff of Q15 (2). De novo assembly was performed on samples using SPAdes version 3.7 (3), and contigs were annotated using Bakta 1.11 (4). Computational pipeline *breseq* was used to identify mutations in the short-read data received from sequencing an evolved clonal isolate relative to the reference sequence NRS6202, NRS6145, or NRS6101 (5) (See Supplemental File F1). As required, genomes were visualised using Artemis 18.1.0 (6) and the browser-based Basic Local Alignment Search Tool (BLAST (7)) was used to compare nucleotide and protein sequences of *comP*. To translate the L6P4 mutated *comP* nucleotide sequence, the translate tool from Expasy was used (8).

#### *Analysis of ComP sequences in Bacillus subtilis genomes sequences*

Operon sequences from the start of ComQ to the end of ComA were extracted from the annotated genome sequences and analysed. Individual CDS/protein sequences were extracted based upon a combination of annotation, and position in the operon relative to the highly conserved ComX. A small number of manual corrections were required to extract the correct sequences in the case of operons with multiple transposon insertions. A concatenated partitioned alignment of 107 essential single-copy genes was generated with bcgTree 1.2.1 (9), including a *Bacillus amyloliquefaciens* sequence (GCA_000772125) as an outgroup. The alignment was used to create a phylogenetic tree using IQ-TREE 3.0.1 (10), using an edge proportional partition model with an 'MF+MERGE' best-fit partition model followed by tree reconstruction using the best partition model, with 1000 ultrafast bootstraps utilising partition resampling with the 'GENESITE' sampling option. Similarly, a multiple sequence alignment of ComQ protein sequences was produced using MUSCLE 5.1 (11) with default parameters, and a phylogenetic tree again produced using IQ-TREE with model selection using ModelFinder Plus (12). Phylogenetic trees were subsequently visualised using iTOL 7.2.1 (13).

*RNA extraction and sequencing data processing*

Biofilms were scraped into tubes into RNAse-free 1.5 ml Eppendorf tubes, and 1 ml stabilization mix (RNAprotect Bacteria Reagent diluted with PBS 2:1 ratio) was added and briefly vortexed. The mix was homogenised using a 23G needle and 2 ml syringe to shear and lyse the cells. Samples were briefly vortexed again to resuspend in the RNAprotect and incubated for 5 minutes at room temperature. 300 μl of the homogenate was then transferred into a new 1.5 ml tube and centrifuged for 10 minutes at 5,000×g at room temperature. 220 μl of a mix containing 200 μl of TE buffer, 3 mg of lysozyme and 20 μl of Proteinase K were added. The tubes were incubated for 30 minutes at 25°C and 550 rpm on a thermomixer. 700 μl of RLT buffer was added into tubes, vortexed for 10 seconds, and the suspension was transferred into a 2 ml tube containing 50 mg of 0.5 mm Zirconia glass beads. The bacterial cells were disrupted by shaking for 10 minutes on the vortex bead adaptor. After homogenization, the tubes were centrifuged for 15 s at 13,000×g and 760 μl of the supernatant was transferred into a new 2-ml tube. 300 μl of chloroform was added, followed by a vigorous shaking by hand for 15 seconds. Tubes were incubated for 10 minutes at RT and then centrifuged for 15 minutes at 13,000×g and 4°C. The upper phase was gently removed into new 1.5 ml tubes, and 590 μl of 80% (v/v) ethanol was added. The suspension was gently mixed by pipetting and 700 μl of it was transferred into a mini spin column and centrifuged for 15 seconds at 13,000×g. This step was repeated until the total volume of suspension was filtered through the mini spin column. The columns were washed in three steps using 700 μl of RW1 buffer (Qiagen) and two times 500 μl or RPE buffer (Qiagen) with centrifugation for 15 seconds at 13,000×g. After the last washing step, the columns were centrifuged for 2 minutes at 13,000×g to dry. The columns were then transferred into new 1.5 ml tubes. 50 uL of RNase-Free water (Qiagen) was applied directly on the column filter and incubated for 2 minutes at RT. The columns were centrifuged for 1 m at 13,000×g. The elution was pipetted onto the column filter again, incubated for a further 2 minutes at RT, and centrifuged for 1 m at 13,000×g. Eluted RNA was transferred to a new 1.5 ml tube. For a DNase treatment, 10 μl of RDD buffer (Qiagen), 2.5 μl of DNase I (Qiagen), and 37.5 μl of RNase-Free water were added and incubated for 10 m at room temperature. 50 μl of 7.5 M LiCl solution was then added, and the tubes were incubated for 1 hour at −20°C. After incubation, the tubes were centrifuged for

15 minutes at 13,000×g and 4°C, the supernatant was discarded and 150 µl of 80% (v/v) ethanol was added. After another centrifugation step for 15 minutes at 13,000×g and 4°C, the supernatant was discarded, and samples were resuspended in 20 µl of RNase-Free water. RNA concentration was estimated using a Nanodrop, and RNA quality was tested by running 500 ng on a 0.8% (w/v) agarose gel run in 1xTAE.

Raw sequencing data were processed to remove ribosomal RNAs using *RiboDetector* 0.3.1. *FastQ Screen* 0.16.0 was then applied to confirm the absence of contamination from other species. Reads were aligned to the NCIB 3610 genome assembly ASM205596v1 (NCBI), using GenBank locus ID CP020102.1 for the chromosome and CP020103.1 for the plasmid pBS32. Plasmid genes were annotated following (1). Non-coding genes were annotated based on sequences extracted from SubtiWiki, which were subsequently mapped to the NCBI genome. A merged GTF file combining NCBI annotations with these non-coding genes is available from (14)]. As NCBI gene symbols differ from those in SubtiWiki (15) (preferred in this study), SubtiWiki genes were mapped in the same way as non-coding sequences, and mismatched gene symbols were altered accordingly.

Read alignment was performed with *STAR* 2.7.11b using the parameters: --sjdbGTFfeatureExon gene --sjdbGTFtagExonParentTranscript gene_id --outFilterType BySJout --outSAMtype BAM SortedByCoordinate --outFilterMultimapNmax 2 --alignIntronMax 1 --limitBAMsortRAM 3000000000 --readFilesCommand zcat --quantMode GeneCounts. The resulting raw counts were imported into Positron and analysed in R using *targets* (16) 1.11.3 pipeline.

Only genes with at least 10 reads in at least one sample were used for differential expression. All samples were included. Differential expression between mutant and control conditions was carried out using *edgeR* 4.6.3. P-values were adjusted for multiple testing with the Benjamini–Hochberg method, and significance was defined as FDR < 0.01 and |logFC| > 1. Functional enrichment was calculated using *fgsea* 1.34.2 with annotations from GO and KEGG, with genes ranked according to -log(FC) log(P), where FC is the fold change and P is the raw, uncorrected p-value from differential expression. The resulting p-values were corrected for multiple tests with the Benjamini–Hochberg method and the significant terms/pathways were defined with FDR < 0.05.

*Confocal Analysis of Colony Biofilms*

Confocal imaging was done using methodology similar to that described previously (17). Mixed isolate colony biofilms were set up by growing the cultures to an $OD_{600}$ of 1 and mixing the strains of interest at a 1:1 ratio as described above. 1 µl of the mixed culture was inoculated on LabTek II 2-chamber microscope vessels (Thermo Scientific), containing 4.4 ml of MSgg agar, solidified and air dried in a laminar flow for 45 minutes. Two biofilms were inoculated per chamber. Imaging was performed using a Leica SP8 upright multiphoton microscope with a chamber pre-warmed to 30 °C. For visualisation of strains producing GFP, the 488 nm argon laser was set to 2% power. For visualisation of mKate2-producing cells multi-photon imaging was performed and the tuneable laser was set to 712 nm at 10% power. The spectral detector was set to collect from 500 nm to 550 nm for the GFP channel and from 600 nm to 690 nm for the mKate2 channel. A resonant mirror at 8 kHz with bi-directional scanning and line averaging of 16x was used and the pinhole was set to 1 AU. The objective used was a 10x, 0.3 NA long working distance dry objective. To image the live biofilms, the lids of the LabTek II 2-chamber vessels were replaced with a #1.5 coverglass to prevent dehydration and shrinking of the agar due to exposure to the flowing air of the microscope chamber. Imaging files were imported, and figures were constructed using OMERO (18).

*Experimental evolution*

Strains NRS6936, NRS6951, and NRS7201 (NRS6105, NRS6145 and NRS6202 respectively adapted to express mTagBFP and containing spectinomycin resistant) and NRS1473 (NCIB 3610 expressing GFP, kanamycin resistant), were used for the experimental evolution. The strains were streaked out on LB plates and incubated at 37°C overnight. The following morning, a day culture of each strain was set up from a single colony in 3 ml of LB and incubated at 37°C with agitation. The cultures were normalised to an $OD_{600}$ of 1 and mixed at a 1:1 ratio. A 5 µl drop of the mixed culture was spotted on MSgg agar, along with a 5 µl drop of each of the single isolate cultures. The plate was incubated at 30°C for 24 hours and imaged. The mixed biofilm was removed from the surface of the MSgg plate with a sterile loop and disrupted by repeatedly passing the material through a 27-gauge in 500 µl of 1x PBS. The sample was serially diluted in PBS buffer and plated onto LB plates containing spectinomycin to select for clones of the strain being adapted. The plates were incubated at 37°C overnight. NRS1473 was also streaked out fresh on LB agar and incubated at 37°C overnight. The following morning, 10 single colonies were picked from the spectinomycin plates, and each one was used to inoculate a 3 ml LB culture. Each of these colonies was considered the first passage of a lineage of evolving clones of the strain being adapted. A 3 ml LB culture of NRS1473 was also set up and all cultures were incubated at 37°C with agitation. The cultures were normalised to an $OD_{600}$ of 1 before each of the cultures of the clones picked from the spectinomycin plates were mixed at a 1:1 ratio with the normalised NRS1473 culture. 1 ml of each of the 10 cultures of the evolving strain was mixed with 800 µl of 50% (v/v) glycerol in cryotubes and stored at -80°C. The mixed cultures along with the single isolate controls were spotted (5 µl drops) on MSgg agar plates. The plates were incubated overnight at 30°C. The following day, the biofilms were imaged and a mixed biofilm containing each of the 10 evolving lineages was removed from the plate, disrupted, serially diluted, and spread over the surface of LB plates containing spectinomycin as described above. NRS1473 was also streaked out on an LB plate. All plates were incubated at 37°C overnight. The following morning, a single colony of each lineage of evolving strain was used to set up a day culture in 3 ml of LB to identify clonal populations that displayed an advantage in coculture compared with the parental population. These clones were considered passage 2. A 3 ml LB culture of NRS1473 was also set up. All cultures were grown at 37°C with agitation and normalised to an $OD_{600}$ of 1. The process of setting up the mixed biofilms, stocking the cultures of evolving lineages, growing biofilms, imaging and so on was repeated as described above. The experiment was stopped after 7 passages. At this point, clones of interest were purified by streaking out to generate single colonies twice before re-stocking them.

#### *RapP and LXG protein sequence analysis*

Proteins in the pfam family PF04740 LXG domain of WXG superfamily (19) were identified in each strain using HMMER (20). A multiple sequence alignment of protein sequences was produced using MUSCLE 5.3 (11) with default parameters and used to construct a phylogenetic tree using IQTree 3.0.1 (10) with a VT+F+I+R5 substitution model selected by ModelFinderPlus (12) with 1000 bootstrap iterations using UFBoot (21). The browser-based Basic Local Alignment Search Tool (BLAST (7)) was used to determine if the protein sequence of RapA, RapB, RapC, RapD, RapE, RapF, RapG, RapH, RapI, RapG, RapK, RapP (as present in NCIB 3610) were present in the genomes of NRS6105, NRS6149, NRS6202. Rstudio was used to generate a heatmap (package pheatmap (10.32614/CRAN.package.pheatmap) using the BLAST results for the sequence with the highest identity (<95%) to show presence and absence.

#### *Structural modelling of ComP*

The protein sequences of ComP from strain NRS6145 (wild type, WT) and the mutant variant carrying the E510K substitution were submitted to the AlphaFold2 server (22) (see Supplemental File F2, Table S8). Predictions were performed in multimer (dimeric) mode, as ComP is known to function as a homodimeric histidine kinase (15, 23). The search space included ADP as a ligand template, to guide modelling of the catalytic ATP-binding domain. For each sequence, five structural models were generated, ranked by AlphaFold confidence scores (pLDDT and predicted alignment error). For each construct, five models were produced and subsequently superimposed. The average pairwise backbone RMSD values within each set were 3.2 Å for the WT and 5.7 Å for the E510K mutant, indicating consistent structural convergence.

Resulting model ensembles for WT and E510K were imported into PyMOL v3.1.0 (Schrödinger, LLC) for structural visualisation and analysis. Models were superimposed based on backbone Cα atoms to assess structural convergence, and root-mean-square deviations (RMSD) were calculated for each ensemble. A global comparison confirmed that both sets of models adopt the characteristic histidine kinase fold previously described for ComP and related proteins (15). The domain organization is conserved, in agreement with published crystallographic and homology-based analyses of histidine kinases (24). Representative models for WT and E510K were selected based on the geometry of residues surrounding position 510. To evaluate the potential local consequences of the substitution, one representative WT and one mutant model were selected based on the orientation of residues surrounding position 510. These models were superposed using the Cα atom of residue 510 as a reference point. Key residues (E510/K510, R331, E394) were inspected and distances between potential hydrogen-bond donor and acceptor atoms were measured, with interactions between 2.5 - 3.5 Å were considered potential hydrogen bonds.

#### *Mathematical model of dual-isolate biofilms*

We used a mathematical model to test hypotheses on the impact of an isolate's doubling time on its competitiveness during biofilm growth. Adapted from an existing mathematical model (25), the model describes the spatio-temporal dynamics of two isolates in a single growing biofilm (Supplemental Methods). In the model, $B_1(x,t)$ and $B_2(x,t)$ denote the respective densities of two isolates at time $t \geq 0$, and space location $x \in \Omega \subset \mathbb{R}^2$, where the space domain $\Omega$ represents the surface of the growth medium in a petri dish. For simplicity, their dynamics are reduced to processes of local growth and spatial spread through density-dependent diffusion and are described by the pair of nondimensionalised partial differential equations

$$\begin{aligned} \frac{\partial \boldsymbol{B_1}}{\partial \boldsymbol{t}} &= \boldsymbol{\nabla} \cdot \left( \boldsymbol{Id}\left(\boldsymbol{1} - \frac{\boldsymbol{B_1} + \boldsymbol{B_2}}{\boldsymbol{k}}\right) \boldsymbol{\nabla B_1} \right) + \boldsymbol{B_1}\left(\boldsymbol{1} - \frac{\boldsymbol{B_1} + \boldsymbol{B_2}}{\boldsymbol{k}}\right), \\ \frac{\partial \boldsymbol{B_2}}{\partial \boldsymbol{t}} &= \boldsymbol{\nabla} \cdot \left( \boldsymbol{Id}\left(\boldsymbol{1} - \frac{\boldsymbol{B_1} + \boldsymbol{B_2}}{\boldsymbol{k}}\right) \boldsymbol{\nabla B_2} \right) + \boldsymbol{rB_2}\left(\boldsymbol{1} - \frac{\boldsymbol{B_1} + \boldsymbol{B_2}}{\boldsymbol{k}}\right). \end{aligned} \tag{1}$$

The indicator function $Id = 1$ if $B_1 + B_2 \leq k$ and $Id = 0$ otherwise, where $k$ is the carrying capacity of the biofilm. The constant $r > 0$ describes the growth rate of isolate $B_2$ during the exponential growth phase relative to that of $B_1$. Due to the nondimensionalisation, it also quantifies the fold difference of the doubling time of $B_1$ during the exponential growth phase to that of $B_2$. At time $t = 0$, the model is initialised by randomly distributing a number of monoclonal subcolonies within the centre of the space domain. Based on CFU counts in biofilm inoculum *vs* CFU counts in mature biofilms (Figure 3C), we set the strain densities to $1/10^5$ of the carrying capacity in these monoclonal subcolonies. This is an improvement to the model compared to a previous application (25). When comparing model outcomes for different values of $r$, we always used the same initial condition to prevent any impact of the random placing of the monoclonal subcolonies. The model was implemented using Matlab's PDE Toolbox. We summarised model results by reporting the relative densities of both strains in percentages at the end of the simulation, that is:

$$\%\, B_1 \text{remaining} = \frac{B_1}{B_1 + B_2}$$

and

$$\%\, B_2 \text{remaining} = \frac{B_2}{B_1 + B_2}$$

respectively.

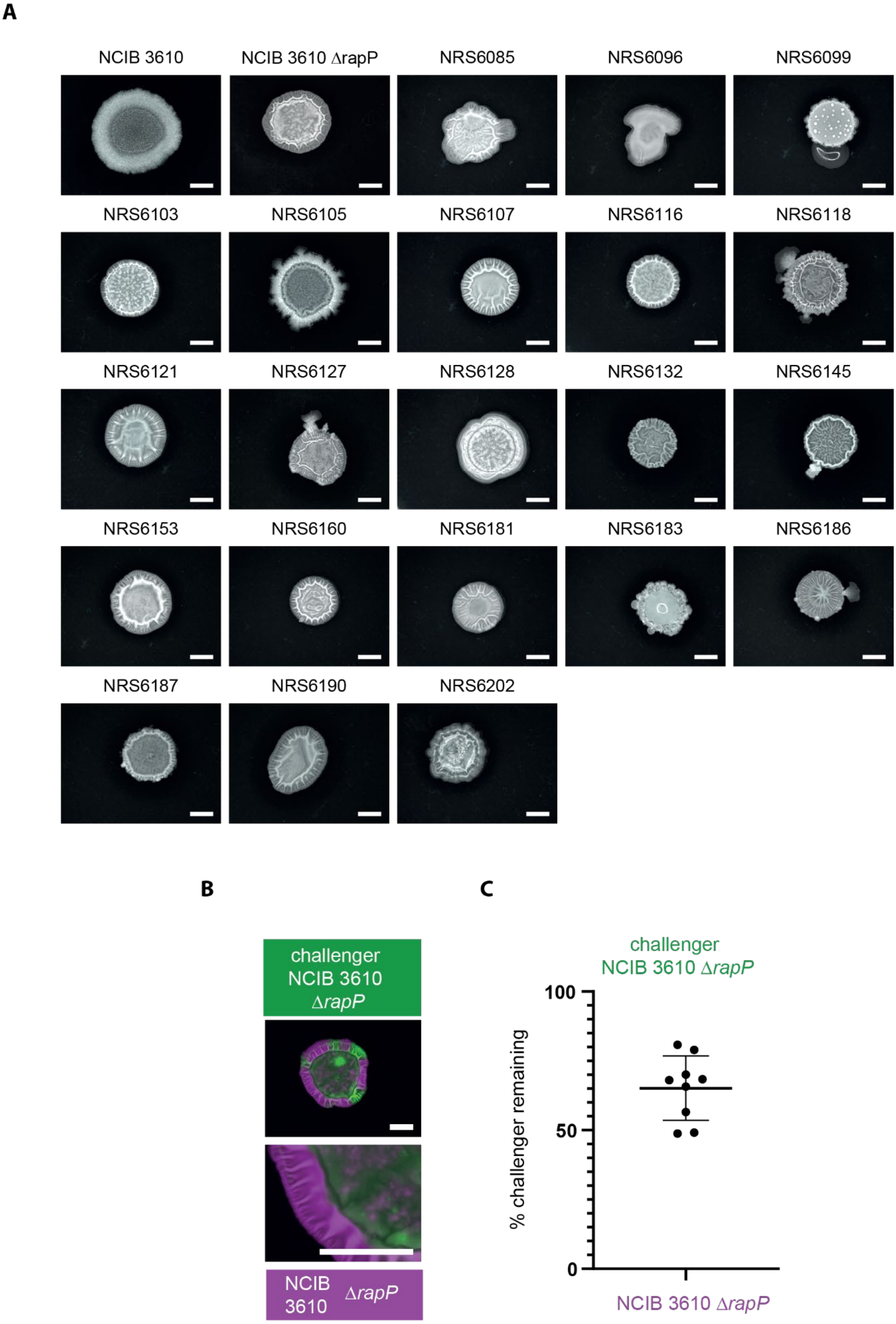
A
NCIB 3610
NCIB 3610 ΔrapP
NRS6085
NRS6096
NRS6099
NRS6103
NRS6105
NRS6107
NRS6116
NRS6118
NRS6121
NRS6127
NRS6128
NRS6132
NRS6145
NRS6153
NRS6160
NRS6181
NRS6183
NRS6186
NRS6187
NRS6190
NRS6202
B
challenger NCIB 3610 ΔrapP
NCIB 3610 ΔrapP
C
challenger NCIB 3610 ΔrapP
100
50
0
% challenger remaining
NCIB 3610 ΔrapP

**Figure S1 Removal of *rapP* reverts NCIB 3610 colony biofilm architecture to that of soil isolates**.

(A) Colony biofilm morphology of NCIB 3610 Δ*rapP* and other soil isolates of *B. subtilis*. Strains were grown on biofilm-inducing media for 48h at 30$^o$C before imaging. The strain and genotype are indicated. The scale bars represent 0.5 cm. All soil isolates shown here are variants constitutively expressing GFP and the image of NCIB 3610 *rapP* is of an mTagBFP-expressing variant.

(B) Representative outcome after 48 hours incubation in a dual isolate colony biofilm incubated at 30$^o$C for 48 hours.

(C) Quantification of the interaction outcome represented as %challenger remaining. The cocultured strain is indicated on the x-axis. Each data point represents a single colony biofilm and is derived from a combination of technical and biological repeats. The error bars represent the standard deviation of the mean.

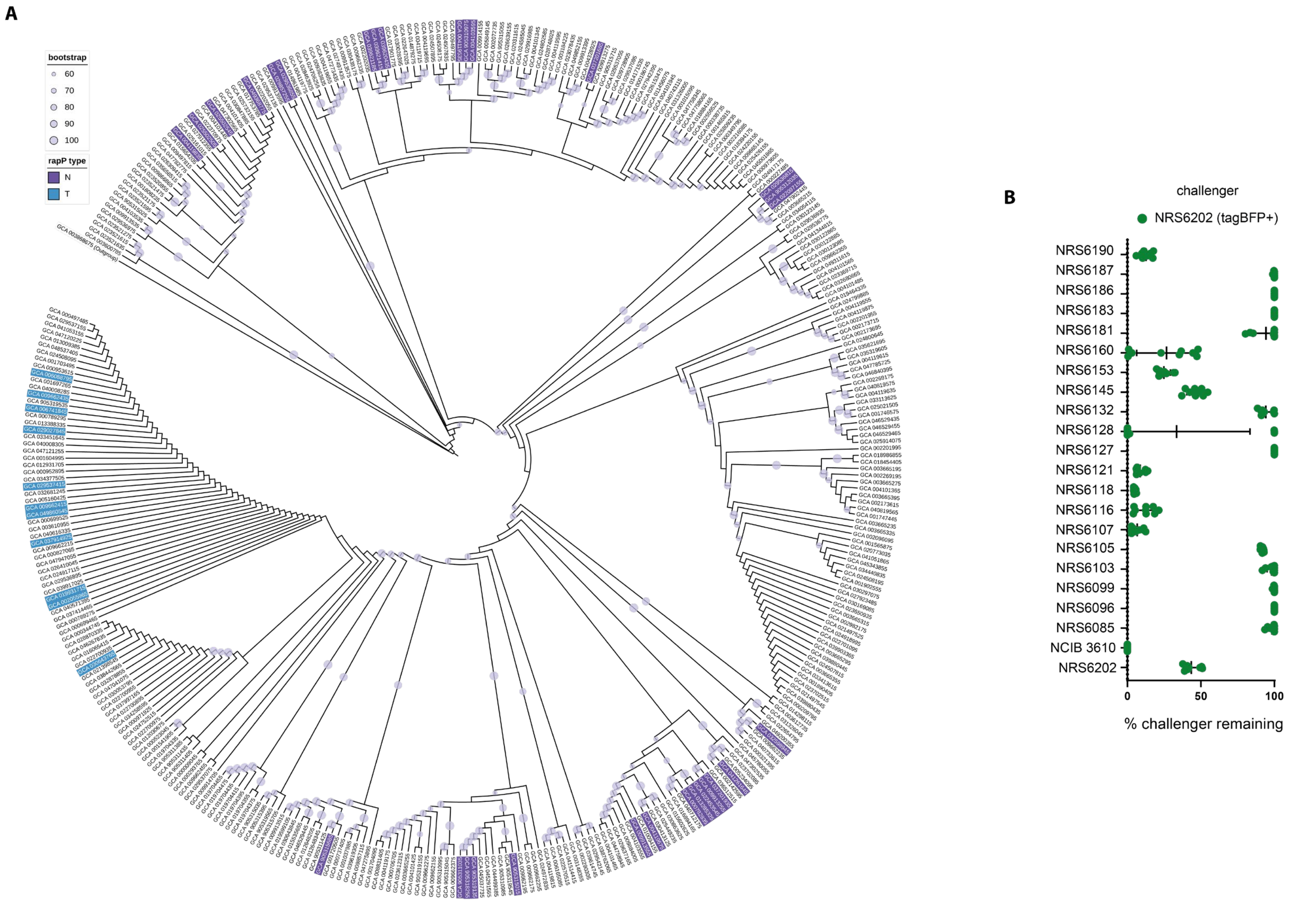
A
bootstrap
60
70
80
90
100
rapP type
N
T
B
challenger
NRS6202 (tagBFP+)
NRS6190
NRS6187
NRS6186
NRS6183
NRS6181
NRS6160
NRS6153
NRS6145
NRS6132
NRS6128
NRS6127
NRS6121
NRS6118
NRS6116
NRS6107
NRS6105
NRS6103
NRS6099
NRS6096
NRS6085
NCIB 3610
NRS6202
0
50
100
% challenger remaining

**Figure S2 – Occurrence of *rapP* and interaction outcome after pairwise screen**

(A) Prevalence of the *rapP* coding region in the genomes of 370 *Bacillus subtilis* isolates. *B. subtilis* isolates indicating isolates carrying *rapP*, and the state of the N236T residue associated with reduced signal responsiveness

(B) Quantification of pairwise biofilm competitions shown as % challenger remaining. Challenger was NRS6202 (mTagBFP+). Partner strains are indicated on the y-axis. Each point = one biofilm; error bars=SD.

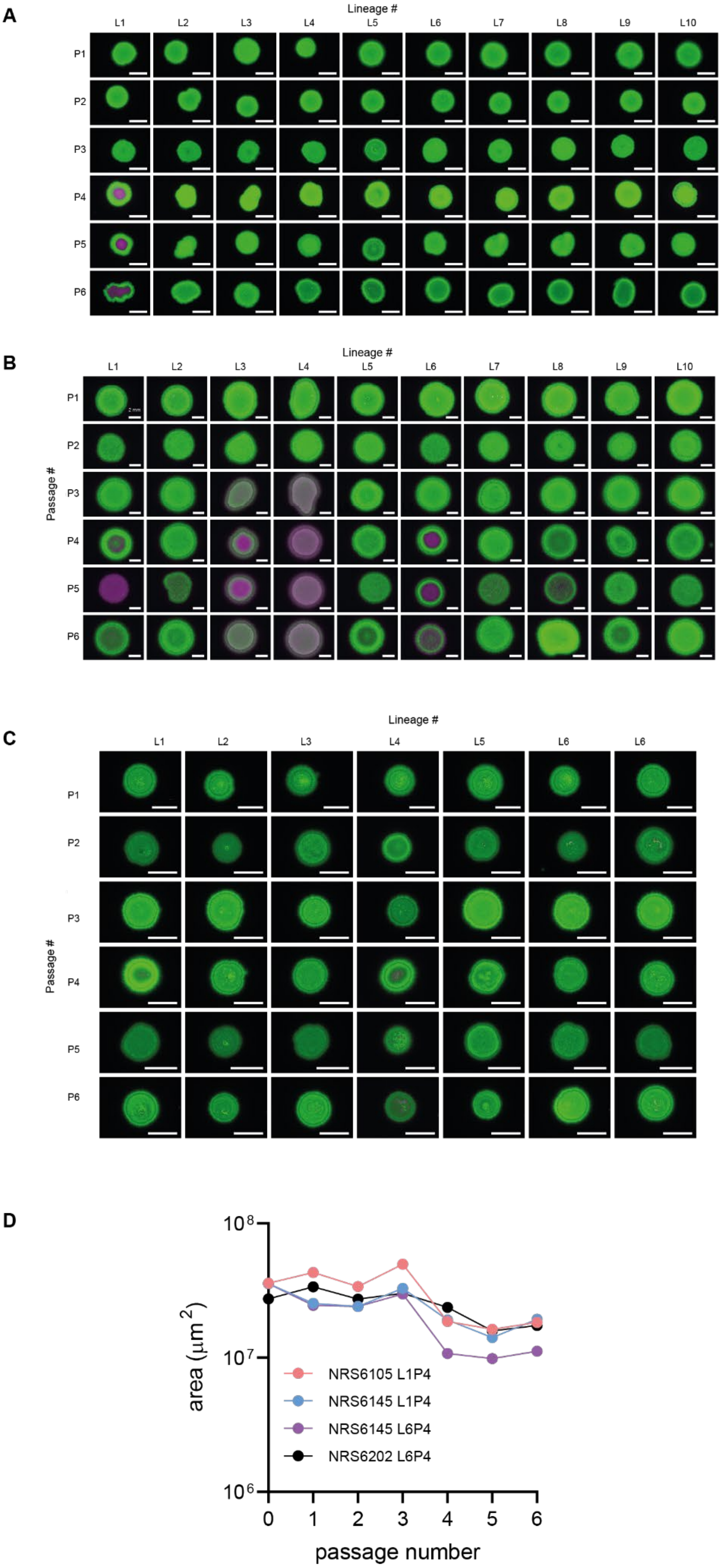
A
Lineage #
L1 L2 L3 L4 L5 L6 L7 L8 L9 L10
P1 P2 P3 P4 P5 P6
B
Lineage #
L1 L2 L3 L4 L5 L6 L7 L8 L9 L10
Passage #
P1 P2 P3 P4 P5 P6
C
Lineage #
L1 L2 L3 L4 L5 L6 L6
Passage #
P1 P2 P3 P4 P5 P6
D
area (μm²)
$10^8$
$10^7$
$10^6$
0 1 2 3 4 5 6
passage number
NRS6105 L1P4
NRS6145 L1P4
NRS6145 L6P4
NRS6202 L6P4

E

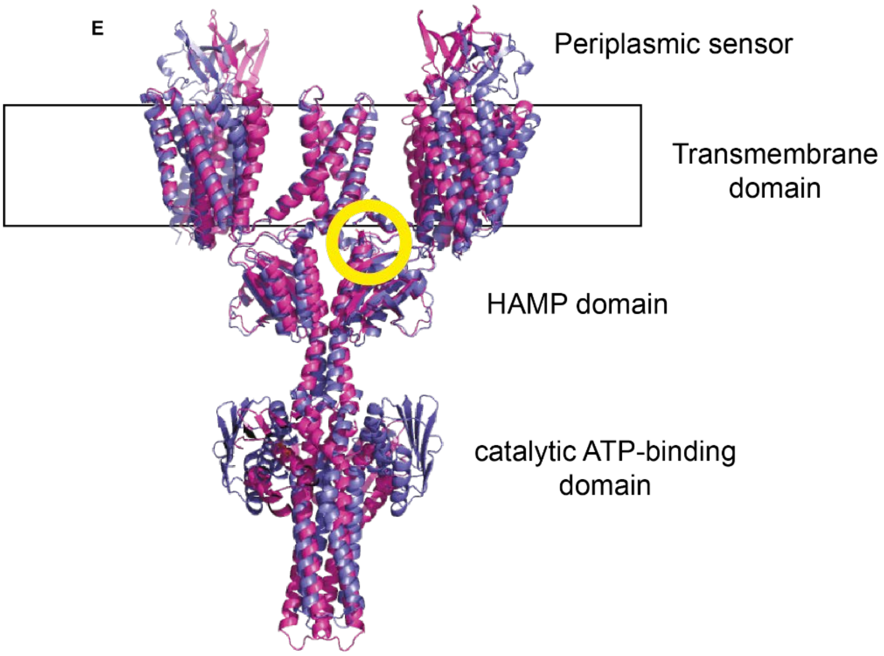
Periplasmic sensor
Transmembrane domain
HAMP domain
catalytic ATP-binding domain

F

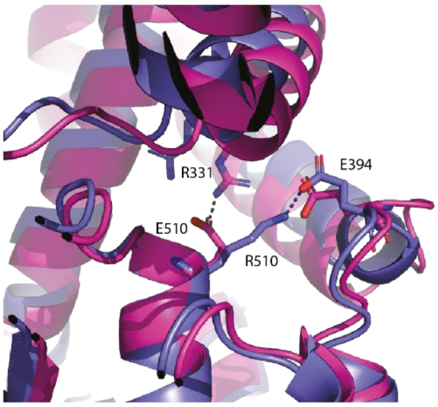
R331
E394
E510
R510

G

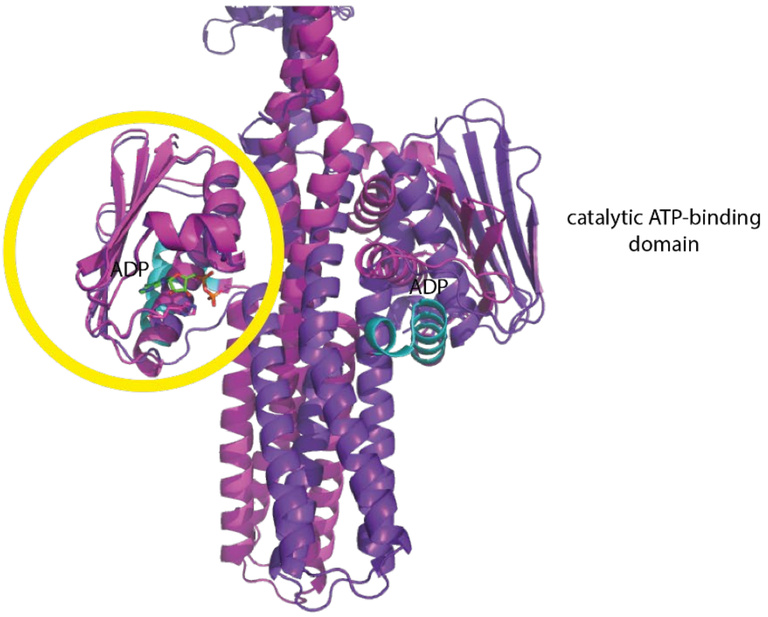


H

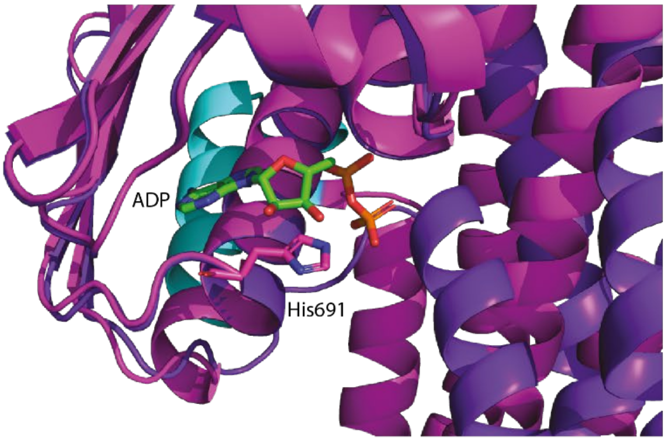


**Figure S3: Lineages of *Bacillus subtilis* subjected to laboratory evolution**

(A-C) Colony biofilms of the evolving soil isolates clones mixed with NCIB 3610 grown on MSgg agar plates for 24h at 30°C before imaging. “mixed” shows the NCIB 3610 (green) and soil isolate (magenta) co-culture biofilms. Panel A shows lineages for NRS6105; panel B shows lineages for NRS6145; panel C shows lineages for NRS6202. P1-P6 on the left-hand side indicates the passage in the experimental evolution experiment and the soil isolate background and lineage shown in each panel are indicated at the bottom, where “L” indicates the lineage and “P” the passage number. The scale bars represent 5 mm (A and C) and 2 mm (B).

(D) The area of the colony biofilms of the evolving soil isolates was quantified with respect to passage number (x-axis) (the images in the “single” panel Figure 3). n=1.

(E-F) Structural models of ComP wild type (WT) and E510R mutant.

(E) AlphaFold-predicted models of the ComP dimer in the WT (pink) and E510R (purple) forms were superimposed and displayed as cartoon representations. The transmembrane (TM) helices are shown at the top and boxed to indicate the approximate position of the lipid bilayer. The periplasmic sensor domain lies between the TM helices, while the cytoplasmic portion consists of the HAMP, DHp, and the catalytic ATP-binding domain typical of histidine kinases. The yellow circle marks the location of residue 510 within at the membrane interface.

(F) Close-up view of the membrane interface domain circled in (E). Side chains of key residues are shown as sticks and coloured as in panel A. Residues R331, E394, and position 510 are labelled. In the WT model (pink), E510 forms potential interactions with R331 and possibly E394 on the adjacent helix. In the E510R mutant (purple), the substitution disrupts this local hydrogen bond network, resulting in a rearranged side-chain interaction pattern (dashed lines).

(G-H) Structural prediction of ComP dimer from NRS6105 (Δ633-649) vs NRS6105 WT.

(G) Predicted structural models were superimposed on the Calpha of one of the two ATP binding domains (circle). The WT protein structure is shown in pink and the mutant Δ633-649 in purple. ADP is shown in sticks, and the deleted 633-649 domain is coloured cyan in the WT protein. The second ATP-binding domain does not superpose between the WT and Δ633-649 structure.

(H) Close-up of the ATP-binding domains of the WT and Δ633-649 mutant. ADP and the side chain of His691 are shown in sticks. The overall ATP-binding site remains largely unchanged between the two proteins.

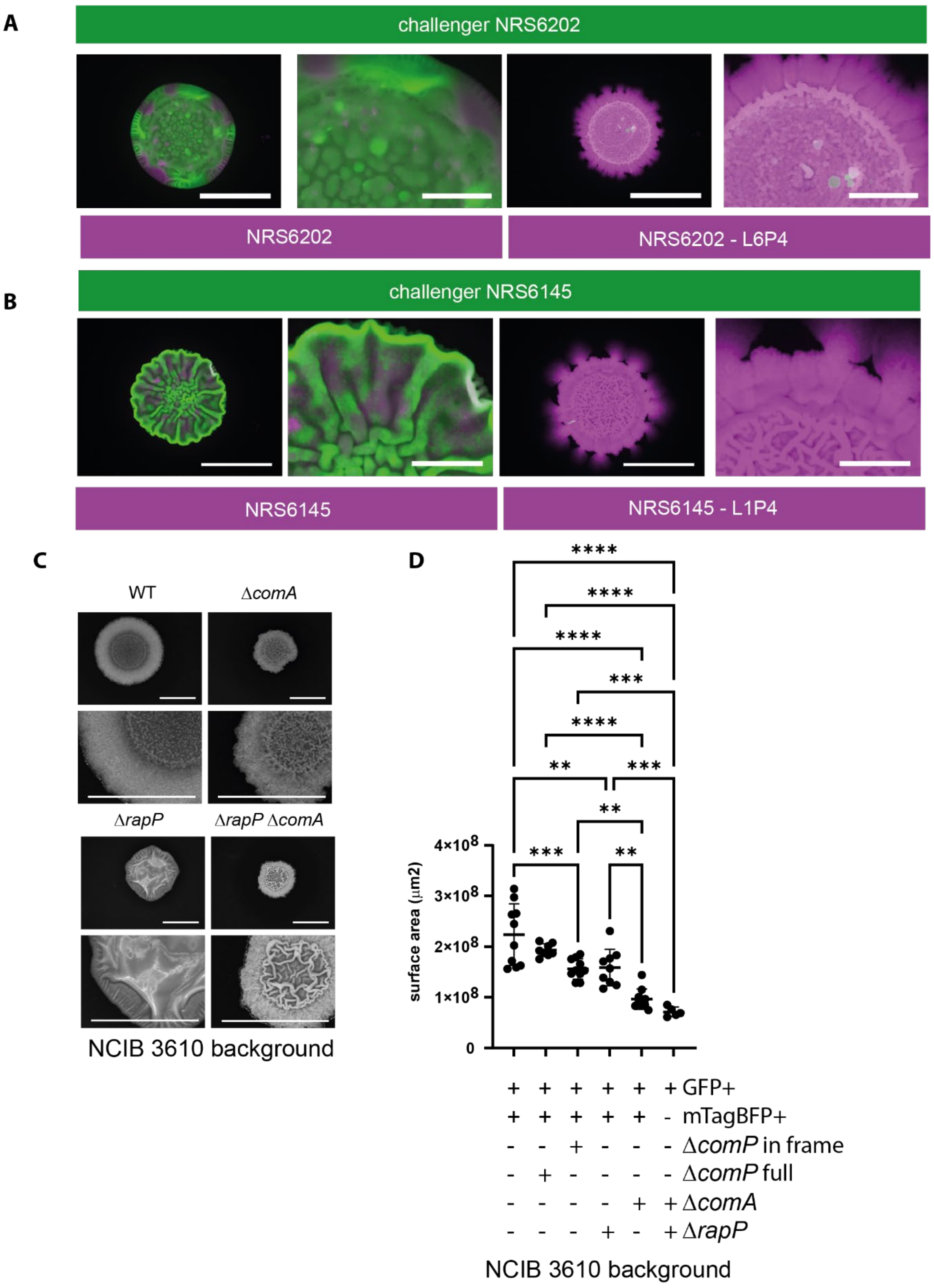


457

458 **Figure S4 ComQXPA influences the outcome of interactions in otherwise**
459 **isogenic contexts.**

(A-B) Representative examples of colony biofilm interactions. The scale bars represent 10 mm or 3 mm. The strains in the coculture are indicated. Images were taken after 48 hours incubation at 30°C.

(C) Colony biofilm morphology after growth on biofilm-inducing media for 48h at 30°C. The strain and genotype are indicated. All strains are GFP+ derivatives of NCIB 3610 (NRS6942, NRS7770, NRS7279 & NRS7771). The scale bars represent 1.0 cm.

(D) Footprint measurements for colony biofilms of the genotypes detailed in the legend of the graph. Ordinary One-way ANOVA Tukey's multiple comparisons test, with a single pooled variance, was used for analysis. Relationships for which no evidence of statistical significance was collected are not shown.  The asterisks represent statistical significance with a p-value of ≤0.05 (*); ≤0.01 (**); ≤0.001 (***) or ≤0.0001 (****).

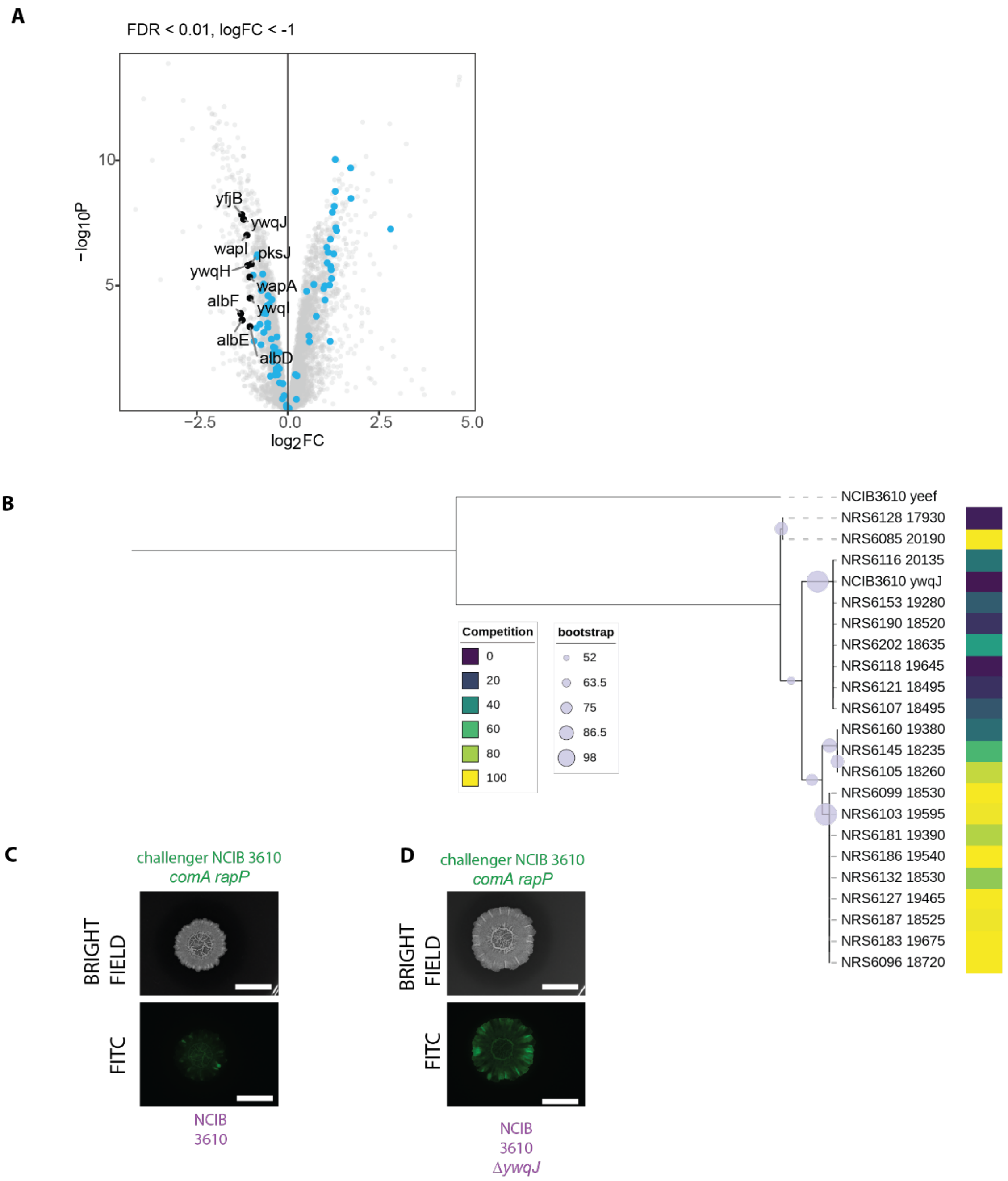


**Figure S5 Identification of a specific competition determinant**

(A) Volcano plot of the 469 differentially expressed genes identified by RNA sequencing. The toxins and other antimicrobials that are expressed to a higher level in NCIB 3610 compared to the Δ*rapP* Δ*comA* strain are indicated.

(B) Phylogenetic tree based upon Muscle alignment of the top matched YwqJ LXG containing protein sequence from the suite of *B. subtilis* isolates relative YwqJ encoded by NCIB 3610. The LXG domain containing protein YeeF from NCIB 3610 was used as an outgroup. The heatmap represents the competition outcome of the different wild isolates when cocultured with NCIB 3610 ∆*rapP* which has an equivalent doubling time is shown (data from Figure 2A).

(C) and (D) A representative outcome of a dual isolate colony biofilm morphology after growth on biofilm-inducing media for 48h at 30$^{o}$C. The strains and genotypes are indicated. The upper panel is bright field, and the lower is the FITC signal false coloured green. The scale bars represent 1.0 cm.

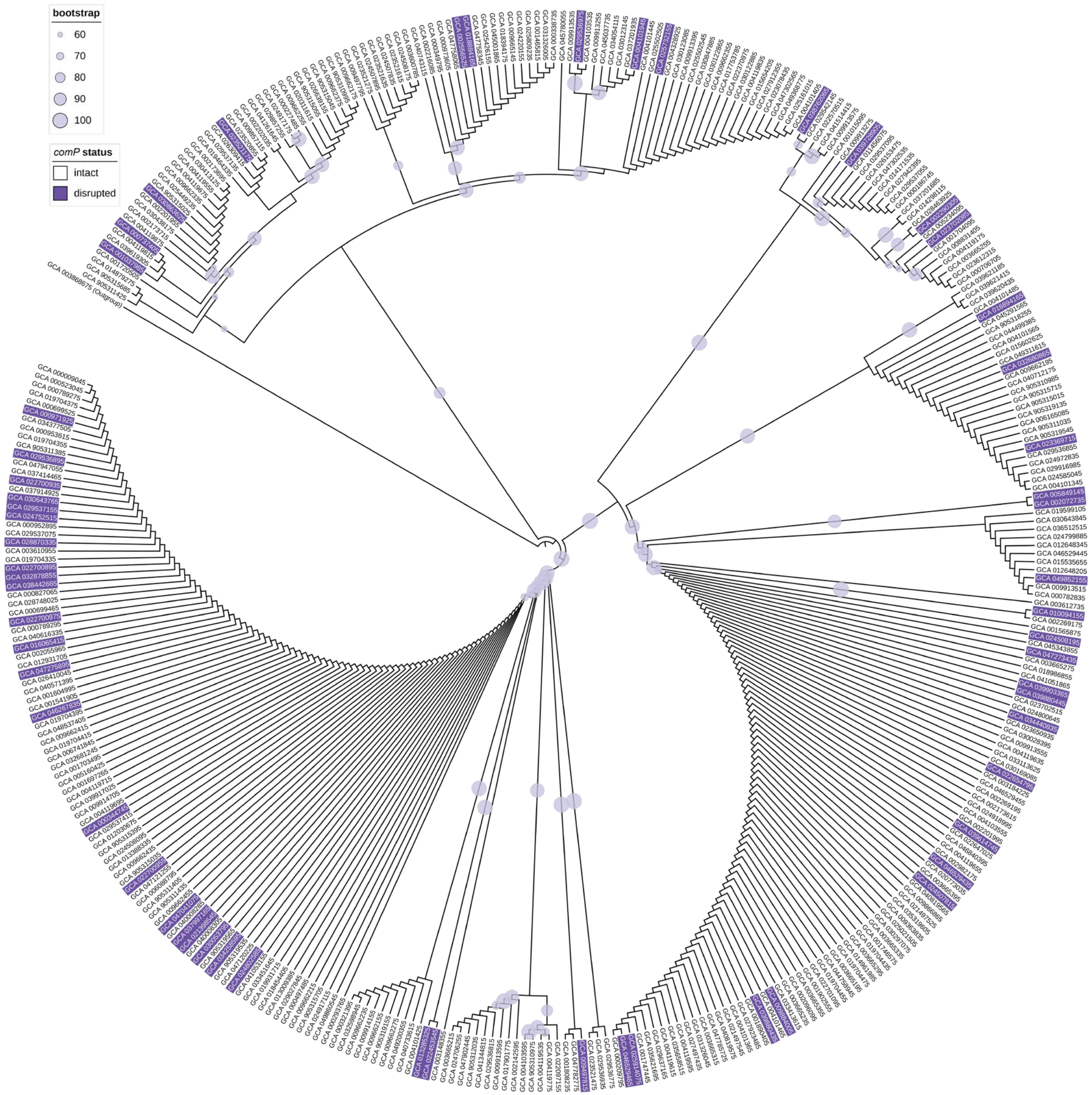
bootstrap
60
70
80
90
100
comP status
intact
disrupted

**Figure S6**

Distribution of disrupted *comP* sequences across phylogeny of 370 *B. subtilis comQ* sequences, selected as a conserved member of the *comQXPA* operon. The phylogeny is rooted on an outgroup consisting of a *Bacillus amyloliquefaciens* sequence. Bootstrap values are represented by differently sized circles on the tree branches.

**Supplemental References**


1. M. A. Konkol, K. M. Blair, D. B. Kearns, Plasmid-encoded ComI inhibits competence in the ancestral strain of Bacillus subtilis. *Journal of Bacteriology* 10.1128/JB.00696-13 (2013).
2. A. M. Bolger, M. Lohse, B. Usadel, Trimmomatic: a flexible trimmer for Illumina sequence data. *Bioinformatics* **30**, 2114-2120 (2014).
3. A. Bankevich *et al.*, SPAdes: a new genome assembly algorithm and its applications to single-cell sequencing. *J Comput Biol* **19**, 455-477 (2012).
4. O. Schwengers *et al.*, Bakta: rapid and standardized annotation of bacterial genomes via alignment-free sequence identification. *Microb Genom* **7** (2021).
5. D. E. Deatherage, J. E. Barrick, Identification of mutations in laboratory-evolved microbes from next-generation sequencing data using breseq. *Methods Mol Biol* **1151**, 165-188 (2014).
6. K. Rutherford *et al.*, Artemis: sequence visualization and annotation. *Bioinformatics* **16**, 944-945 (2000).
7. E. W. Sayers *et al.*, Database resources of the National Center for Biotechnology Information in 2025. *Nucleic Acids Res* **53**, D20-D29 (2025).
8. E. Gasteiger *et al.*, ExPASy: The proteomics server for in-depth protein knowledge and analysis. *Nucleic Acids Res* **31**, 3784-3788 (2003).
9. M. J. Ankenbrand, A. Keller, bcgTree: automatized phylogenetic tree building from bacterial core genomes. *Genome* **59**, 783-791 (2016).
10. B. Q. Minh *et al.*, IQ-TREE 2: New Models and Efficient Methods for Phylogenetic Inference in the Genomic Era. *Mol Biol Evol* **37**, 1530-1534 (2020).
11. R. C. Edgar, MUSCLE: multiple sequence alignment with high accuracy and high throughput. *Nucleic Acids Res* **32**, 1792-1797 (2004).
12. S. Kalyaanamoorthy, B. Q. Minh, T. K. F. Wong, A. von Haeseler, L. S. Jermiin, ModelFinder: fast model selection for accurate phylogenetic estimates. *Nat Methods* **14**, 587-589 (2017).
13. I. Letunic, P. Bork, Interactive Tree of Life (iTOL) v6: recent updates to the phylogenetic tree display and annotation tool. *Nucleic Acids Res* **52**, W78-W82 (2024).
14. N. Stanley-Wall, J. C. Abbott, Regulatory-Rewiring-Drives-Intraspecies-Competition-in-Bacillus-subtilis - Software. Zendod.
15. T. Pedreira, C. Elfmann, J. Stulke, The current state of SubtiWiki, the database for the model organism Bacillus subtilis. *Nucleic Acids Res* **50**, D875-D882 (2022).
16. W. M. Landau, The targets R package: a dynamic Make-like function-oriented pipeline toolkit for reproducibility and high-performance computing. *Journal of Open Source Software* **6** (2022).
17. M. Porter, F. A. Davidson, C. E. MacPhee, N. R. Stanley-Wall, Systematic microscopical analysis reveals obligate synergy between extracellular matrix components during Bacillus subtilis colony biofilm development. *Biofilm* **4** (2022).
18. C. Allan *et al.*, OMERO: flexible, model-driven data management for experimental biology. *Nat Methods* **9**, 245-253 (2012).

19. T. Paysan-Lafosse *et al.*, InterPro in 2022. *Nucleic Acids Res* **51**, D418-D427 (2023).
20. R. D. Finn, J. Clements, S. R. Eddy, HMMER web server: interactive sequence similarity searching. *Nucleic Acids Res* **39**, W29-37 (2011).
21. D. T. Hoang, O. Chernomor, A. von Haeseler, B. Q. Minh, L. S. Vinh, UFBoot2: Improving the Ultrafast Bootstrap Approximation. *Mol Biol Evol* **35**, 518-522 (2018).
22. J. Abramson *et al.*, Accurate structure prediction of biomolecular interactions with AlphaFold 3. *Nature* **630**, 493-500 (2024).
23. S. Lazaridi, J. Yuan, T. Lemmin, Atomic insights into the signaling landscape of E. coli PhoQ histidine kinase from molecular dynamics simulations. *Scientific reports* **14**, 17659 (2024).
24. F. Jacob-Dubuisson, A. Mechaly, J. M. Betton, R. Antoine, Structural insights into the signalling mechanisms of two-component systems. *Nat Rev Microbiol* **16**, 585-593 (2018).
25. L. Eigentler *et al.*, Founder cell configuration drives competitive outcome within colony biofilms. *ISME J* 10.1038/s41396-022-01198-8 (2022).
